# Meiosis reveals the early steps in the evolution of a neo-XY sex chromosome pair in the African pygmy mouse *Mus minutoides*

**DOI:** 10.1101/2020.06.29.177329

**Authors:** Ana Gil-Fernández, Paul A. Saunders, Marta Martín-Ruiz, Marta Ribagorda, Pablo López-Jiménez, Daniel L. Jeffries, María Teresa Parra, Alberto Viera, Julio S. Rufas, Nicolas Perrin, Frederic Veyrunes, Jesús Page

## Abstract

Sex chromosomes of eutherian mammals are highly different in size and gene content, and share only a small region of homology (pseudoautosomal region, PAR). They are thought to have evolved through an addition-attrition cycle involving the addition of autosomal segments to sex chromosomes and their subsequent differentiation. The events that drive this process are difficult to investigate because sex chromosomes in most mammals are at a very advanced stage of differentiation. Here, we have taken advantage of a recent translocation of an autosome to both sex chromosomes in the African pygmy mouse *Mus minutoides*, which has restored a large segment of homology (neo-PAR). By studying meiotic sex chromosome behavior and identifying fully sex-linked genetic markers in the neo-PAR, we demonstrate that this region shows unequivocal signs of early sex-differentiation. First, synapsis and resolution of DNA damage intermediates are delayed in the neo-PAR during meiosis. Second, recombination is suppressed in a large portion of the neo-PAR. However, the inactivation process that characterizes sex chromosomes during meiosis does not extend to this region. Finally, the sex chromosomes show a dual mechanism of association at metaphase-I that involves the formation of a chiasma in the neo-PAR and the preservation of an ancestral achiasmate mode of association in the non-homologous segments. We show that the study of meiosis is crucial to apprehend the onset of sex chromosome differentiation, as it introduces structural and functional constrains to sex chromosome evolution. Synapsis and DNA repair dynamics are the first processes affected in the incipient differentiation of X and Y chromosomes, and they may be involved in accelerating their evolution. This provides one of the very first reports of early steps in neo-sex chromosome differentiation in mammals, and for the first time a cellular framework for the addition-attrition model of sex chromosome evolution.

**AUTHOR SUMMARY:** The early steps in the evolution of sex chromosomes are particularly difficult to study. Cessation of recombination around the sex-determining locus is thought to initiate the differentiation of sex chromosomes. Several studies have investigated this process from a genetic point of view. However, the cellular context in which recombination arrest occurs has not been considered as an important factor. In this report, we show that meiosis, the cellular division in which pairing and recombination between chromosomes takes place, can affect the incipient differentiation of X and Y chromosomes. Combining cytogenetic and genomic approaches, we found that in the African pygmy mouse *Mus minutoides*, which has recently undergone a sex chromosome-autosome fusion, synapsis and DNA repair dynamics are altered along the newly added region of the sex chromosomes, likely interfering with recombination and thus contributing to the genetic isolation of a large segment of the Y chromosome. Therefore, the cellular events that occur during meiosis are crucial to understand the very early stages of sex chromosome differentiation. This can help to explain why sex chromosomes evolve very fast in some organisms while in others they have barely changed for million years.

## INTRODUCTION

Sex chromosomes in mammals originated when one member of a pair of autosomes acquired a male-determining allele [1]. Nowadays, the X and Y chromosomes in most species are highly differentiated in both size and gene content. Differentiation is thought to have been initiated by a suppression of recombination around the sex-determining locus during male meiosis, followed by an extension of the non-recombining region, possibly involving chromosomal rearrangements and sexually antagonistic selection [2-4]. Recombination suppression, therefore, is a key trigger of sex chromosome evolution, allowing the independent evolution of X and Y chromosomes, and notably causing the degeneration of the Y [5].

The sex chromosomes of all therian mammals (marsupials and eutherians) have a common origin. However, these two groups diverged from each other about 180 million years ago (mya) and their sex chromosomes have reached different points of differentiation [6, 7]. In most marsupials, the sex chromosomes are completely differentiated and no longer share a homologous region [8, 9]. In eutherians, following a period of significant differentiation but prior to the radiation of the group about 117 mya, the translocation of an autosomal segment to both sex chromosomes expanded the homology of the small region that still recombined, i.e., the pseudoautosomal region (PAR) [6, 10]. Afterwards, the Y chromosome engaged in a new cycle of differentiation and homology with the X chromosome was again reduced. However, the pace of this process has varied in different groups. For instance, although most eutherians still conserve a small segment of the PAR, in several groups such as gerbils [11, 12], voles [13-15] and pygmy mice [16], the X and Y chromosomes are completely differentiated (and have no PAR). A few species have even completely lost the Y chromosome, including voles of the genus *Ellobius* [17] and the Ryukyu spiny rat *Tokudaia osimensis* [18]. In contrast, in some bats [19], bovids [20], primates [21] and rodents [16, 22], new autosomal translocations have, once again, restored a large section of the PAR, which can initiate a new differentiation process. This situation reveals a scenario in which sex chromosomes are continuously evolving in a process that has been called the addition-attrition cycle [8, 23].

Recombination suppression, which triggers sex chromosome differentiation, can be caused by different mechanisms (reviewed in [24]. However, it is usually disregarded that recombination occurs during meiosis, a highly specialized cell division during which chromosomes must go through synapsis, recombination and segregation [25]. These complex processes, respectively, involve: 1) the association of homologous chromosomes in pairs (synapsis), through a meiosis-specific structure called the synaptonemal complex (SC); 2) the exchange of genetic information in a process called meiotic recombination, a DNA repair mechanism that leads to the formation of physical connections between homologous chromosomes called chiasmata and 3) the segregation of homologous chromosomes to the daughter cells during first the meiotic division. These processes are interdependent, as synapsis defects leads to recombination disturbance, and vice versa, and defects on either synapsis or recombination have a great impact on chromosome segregation and fertility [26-31]. In most eutherian species, X and Y chromosomes are able to achieve these processes during male meiosis because they have a small PAR whose conserved homology allows the chromosomes to synapse, recombine and form a chiasma [12, 32], thus maintaining an association between them until they segregate during the first meiotic division. Nevertheless, the extreme differentiation of sex chromosomes has consequences on meiotic behavior, as synapsis and reciprocal recombination are not possible in the differentiated (non-homologous) segments, and both processes are conspicuously delayed in the homologous region (PAR) [32, 33]. Moreover, the presence of large unsynapsed chromosomal segments affects the transcriptional activity of sex chromosomes by triggering a silencing process called meiotic sex chromosome inactivation (MSCI) [34-36]. In species with completely differentiated sex chromosomes, synapsis and recombination are absent (sex chromosomes are asynaptic and achiasmate) and segregation is ensured by alternative mechanisms [11, 15, 37-40].

Main clues about the initial steps of sex chromosome differentiation have traditionally come from genetic studies [2, 3, 5, 41]. However, investigating how sex chromosomes stop recombining during meiosis and how meiotic modifications of their behavior arise are crucial to understand the early stages of sex chromosome evolution. This is particularly challenging in mammals, because sex chromosomes in most species are at an advanced stage of divergence, revealing little information about the processes that initiated their differentiation. Hence, less divergent systems that are at earlier stages of evolution are needed to study these processes. Sex chromosome-autosome fusions are ideal candidates to shed light on this process [3, 41, 42]. These fusions are rare in mammals as they tend to cause somatic and reproductive defects [43] by interfering with the process of sex chromosome inactivation [44, 45], among others. Nevertheless, examples of viable fusions are available [19-22, 46].

Here, we report the meiotic behavior of a neo-sex chromosome pair in the African pygmy mouse *Mus minutoides*. Sex chromosomes in most species of pygmy mice are completely differentiated and are thus achiasmate. However, in *M. minutoides*, a fusion of both the X and Y chromosomes with autosomal pair 1 about 0.9-1.6 mya gave rise to a neo-sex chromosome pair with a large neo-PAR [47-49]. After this addition event, the Y chromosome may have entered a new attrition phase, which should affect the behavior of sex chromosomes in male meiosis. To test this hypothesis, we have analyzed synapsis, recombination and segregation of the sex chromosomes in *M. minutoides*. Our results show that the neo-PAR displays signs of modification in the processes of synapsis and DNA repair in male meiosis, suggesting an incipient process of “sexualization”. By combining a cytogenetic and a genotyping-by-sequencing approach (using restriction site-associated DNA sequencing, RAD-seq), we demonstrate that recombination between the X and Y chromosomes has ceased in an extensive portion of the neo-PAR proximal to the centromere. However, MSCI does not extend to this region. Finally, we observed a dual mechanism of association displayed by the sex chromosomes during metaphase-I, which involves a chiasmate and an achiasmate association, a finding that has, to our knowledge, not been previously reported. Overall, these findings illustrate the cellular modifications undergone by mammalian sex chromosomes at early stages of differentiation and offer a framework to understand the subsequent steps in their evolution.

## RESULTS

*Mus minutoides* harbors a particularly variable karyotype owing to the high frequency of Robertsonian (Rb) translocations involving autosomes, with diploid numbers ranging from 2n=18 to 2n=36. This species is also characterized by the presence of a Rb translocation between chromosome 1 and the X and Y chromosomes (Fig. 1A and B) [49]. The population studied here, coming from the Caledon Nature Reserve in South Africa, has a diploid number of 2n=18 and all chromosomes are metacentric. In males, the sex chromosome pair is composed of (i) non-homologous arms, formed by the completely differentiated (achiasmate) “ancestral” sex chromosomes that differ in length, and (ii) homologous arms, formed by ancestral chromosome 1 (hereafter, the neo-PAR) that are homomorphic in length and in their G-banding pattern (Fig. 1A, B) [48]. During meiosis, chromosomes show highly extended centromeres (Fig. 1C and D), a feature that is not present in the Rb chromosomes of other mouse species, including *Mus musculus domesticus* [50], suggesting it is specific to the Rb translocation mechanism in pygmy mice [48, 51, 52]. The extended centromeres, which vary in length, do not assemble any components of the SC, neither the axial/lateral elements (AEs/LEs) nor the transverse filaments (TFs) (supplementary Fig. S1, S2). We confirmed that this result was not an artifact of the spreading technique as it was also observed in spermatocyte squashes (supplementary Fig. S1), which preserve chromatin condensation and organization.

**Figure 1.**
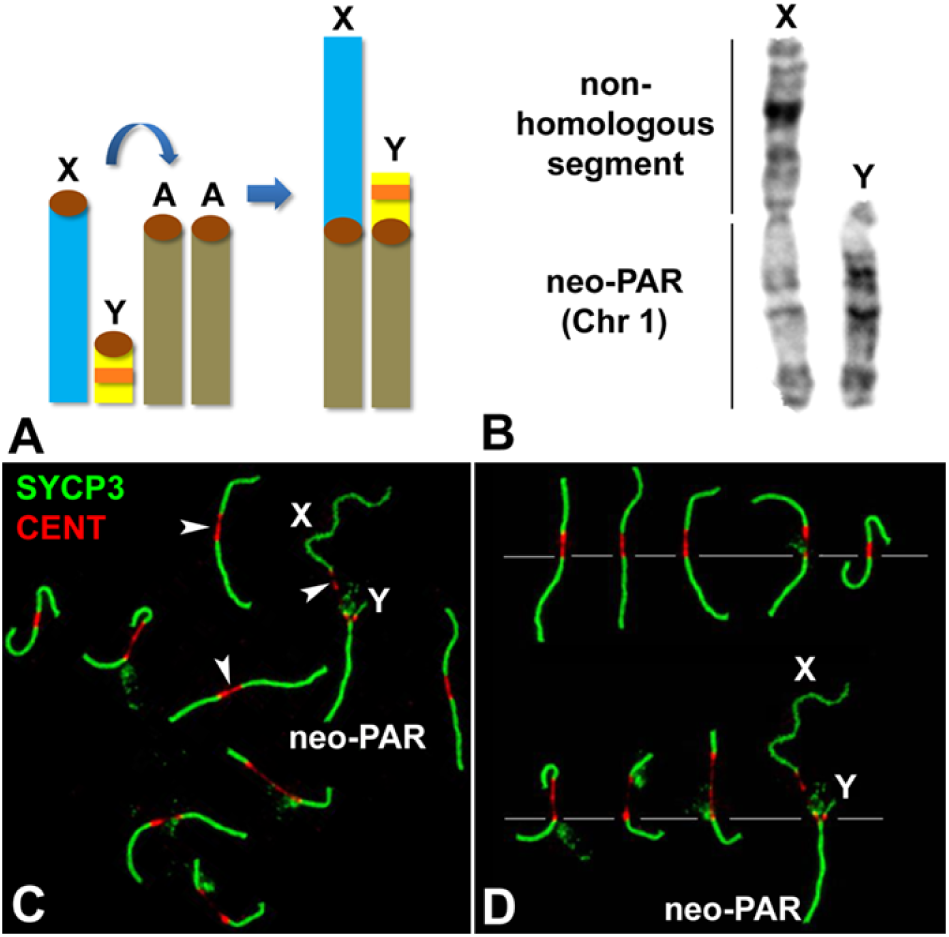
Sex chromosomes in *M. minutoides*. **A**. Schematic representation of the Rb translocation between the sex chromosomes (X and Y) and autosomal chromosome 1 (A). The ancestral sex chromosomes are completely differentiated and, thus, achiasmate. The translocation results in a large neo-PAR constituted by chromosome 1. The orange band in the Y chromosome represents the sex-determining locus. **B**. G-banding of the sex chromosomes indicating the neo-PAR and the non-homologous segment. **C**. Spread spermatocyte at pachytene labelled with antibodies against SYCP3 (green) and centromeres (red). The trajectories of SCs are interrupted at the centromeres, which appear largely stretched (arrowheads). Non-homologous segments of the sex chromosomes (X, Y) and the neo-PAR are indicated. **D**. Meiotic karyotype of *M. minutoides*. Bivalents are arranged according to the length of their largest arm, as previously described [48, 77]. The sex bivalent is largely heteromorphic in the non-homologous segments.

### The neo-PAR shows synapsis and DNA repair delay

To analyze the meiotic behavior of the male sex chromosomes, we first studied the sequence of chromosome synapsis by characterizing the immunolocalization of SYCP3, a marker of the AEs/LEs of the SC, and γH2AX (histone H2AX phosphorylated at serine 139), which marks chromosomal regions that present DNA damage and/or have not completed synapsis [53]. We focused on the specific behavior of the sex chromosomes (the complete cycle of chromosome synapsis during prophase-I is presented in supplementary Fig. S3). X and Y can be unambiguously discerned from the rest of the chromosomes at late zygotene (Fig. 2A). At this stage, the neo-PAR is only partially synapsed. Synapsis in sex chromosomes usually starts at the distal end of the neo-PAR and progresses towards the centromere, while the non-homologous regions (ancestral X and Y) remain separated. The chromatin surrounding the unsynapsed AEs of the sex chromosomes, in both the non-homologous segments and the neo-PAR, are strongly labelled with γH2AX. At the beginning of pachytene (Fig. 2B-B’’), while the autosomes are completely synapsed and present only scattered small γH2AX foci, the region of the neo-PAR proximal to the centromere, comprising about one third of this chromosome arm, appears unsynapsed and broadly labelled with γH2AX. This, clearly indicates that synapsis of the neo-PAR is delayed in relation to autosomes. The unsynapsed region of the neo-PAR tends to decrease as pachytene proceeds (Fig. 2C-C’’), with the neo-PAR finally appearing completely synapsed at late pachytene (Fig. 2D-D’’). We corroborated the assembly of a mature SC in this region by characterizing the localization of SYPC1, the main component of TFs (supplementary Fig. S2). As expected, the non-homologous X and Y segments remain unsynapsed and intensively labelled by γH2AX throughout prophase-I. We never observed synapsis extending to these chromosomal regions, although they appear associated forming a typical sex body.

**Figure 2.**
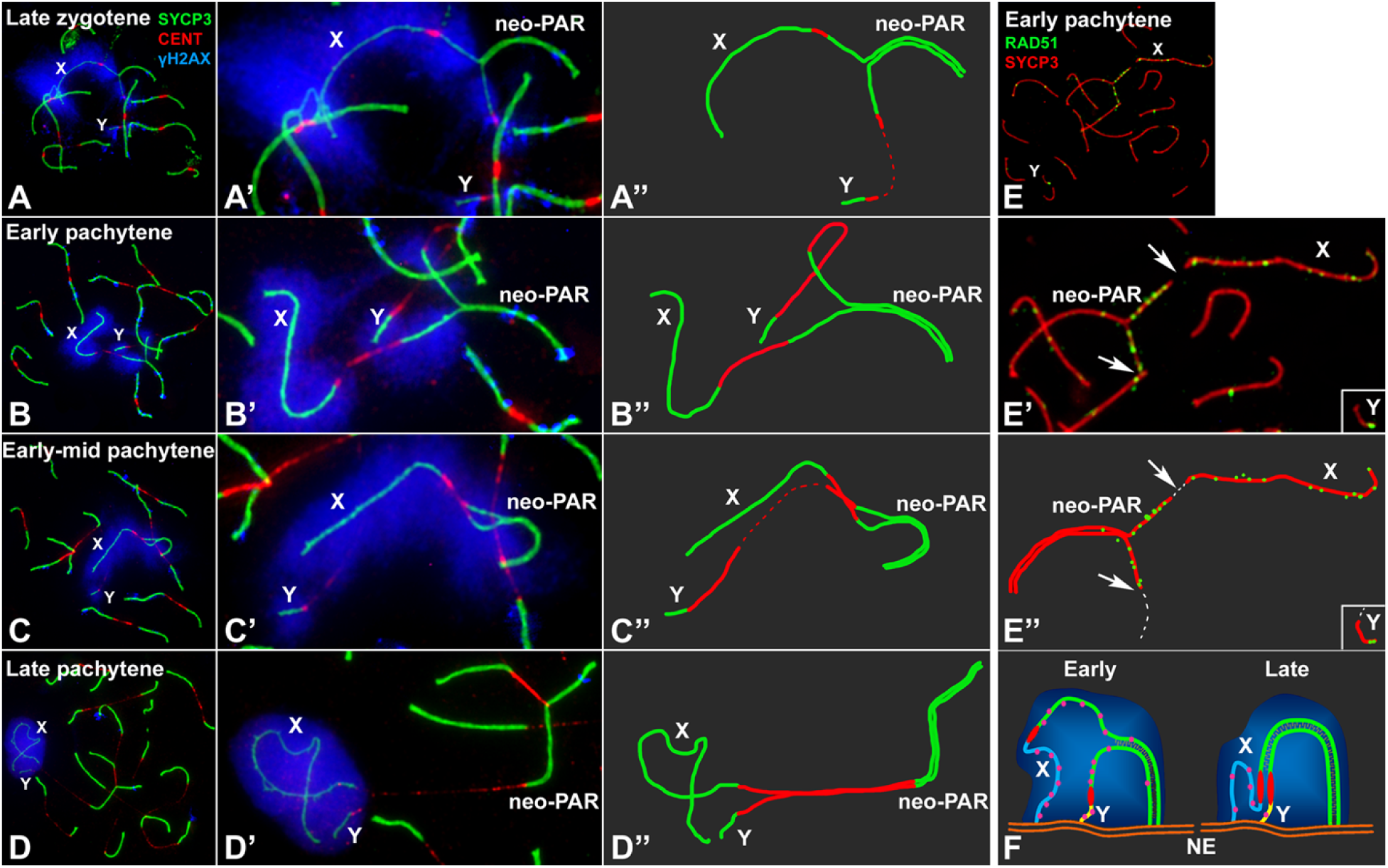
Synapsis and DNA repair in the sex chromosomes. **A-D’’**. Spread spermatocytes in prophase-I labelled with antibodies against SYCP3 (green), centromeres (red) and γH2AX (blue). Non-homologous segments of the sex chromosomes (X, Y) and the neo-PAR are indicated. **A-A’’**. Zygotene. Most autosomes have completed synapsis. Enlarged view of the sex chromosomes (**A’**) and their schematic representation (**A’’**). Both the non-homologous segments and a large portion of the neo-PAR are unsynapsed and labelled with γH2AX. In some cases, centromeres are so elongated that the signal is very weak (represented as dashed lines). **B-B’’**. Early pachytene. Autosomes have fully synapsed but sex chromosomes still have not completed synapsis in the neo-PAR. The centromeres appear greatly stretched. **C-C’’**. Mid-pachytene. Synapsis in the sex chromosomes has extended but still is not complete. **D-D’’**. Late pachytene. The neo-PAR has completely synapsed. Although the non-homologous segments of the sex chromosomes remain unsynapsed, they are in close proximity, forming a single chromatin mass that is strongly labelled with γH2AX. **E-E’’**. Details and schematic representation of a sex bivalent from an early pachytene spermatocyte labelled with antibodies against SYCP3 (red) and RAD51 (green). The proximal regions of the neo-PAR are unsynapsed and accumulate RAD51 foci. The non-homologous segments also accumulate RAD51. Arrows and dashed lines (in E’’) indicate putative centromere positions. **F**. Schematic representation of synapsis in the sex chromosomes. At early pachytene, synapsis proceeds from the distal end of the neo-PAR; however, compared with autosomes, it is delayed. Unsynapsed regions accumulate DNA repair proteins (pink foci). As synapsis proceeds, the X chromosome bends to allow the complete synapsis of the neo-PAR. Repair proteins are also removed from the neo-PAR but remain on the unsynapsed non-homologous segments. NE: nuclear envelope.

We next analyzed the dynamics of DNA repair on the sex chromosomes by characterizing the localization of RAD51, a component of the recombination machinery involved in the interaction between the DNA sequences of homologous chromosomes (Fig. 2E-E’’ and supplementary Fig. S4). RAD51 localizes as small foci on the AEs/LEs where DNA double strand breaks are being repaired. As in other mammals [54, 55], we found that, in *M. minutoides*, RAD51 removal from the sex chromosomes is delayed compared with the autosomes. At early pachytene, RAD51 has mostly disappeared from the autosomes but is still abundant not only on the non-homologous segments of the sex chromosomes but also in the unsynapsed regions of the neo-PAR (Fig. 2E-E’’). These results reveal lagging DNA repair dynamics in the proximal region of the neo-PAR, which could be caused by the synaptic delay and/or by a defective recognition of DNA sequences along this chromosomal segment (Fig. 2F).

### MSCI does not extend to the neo-PAR

Since the presence of γH2AX during pachytene is related with the inactivation of sex chromosomes [36, 53], we wanted to ascertain if the neo-PAR segment that exhibits delayed synapsis is subjected to MSCI. To do this, we assessed the localization of additional MSCI markers (Fig. 3). We first examined the distribution of RNA polymerase-II to determine the level of transcriptional activity (Fig. 3A-D). In line with previous reports [33, 56], a transcription burst is observed from late pachytene onward on all of the chromosomes except the non-homologous segments of the sex chromosomes (Fig. 3C and D). At this time, the neo-PAR has already completed synapsis and, alike autosomes, is labelled by RNA polymerase-II. We also studied the localization of ATR kinase, which is involved in the phosphorylation of γH2AX during MSCI [36, 57]. ATR appears along unsynapsed AEs of autosomes and sex chromosomes at zygotene (Fig. 3E). By mid pachytene, ATR is observed only on the unsynapsed AEs of the sex chromosomes (Fig. 3F). By late pachytene (Fig. 3G) and diplotene (Fig. 3H), ATR localization has extended to the unsynapsed chromatin of only the non-homologous regions of these chromosomes, suggesting that the neo-PAR completes synapsis before the transcription burst and, therefore, is not subjected to MSCI.

**Figure 3.**
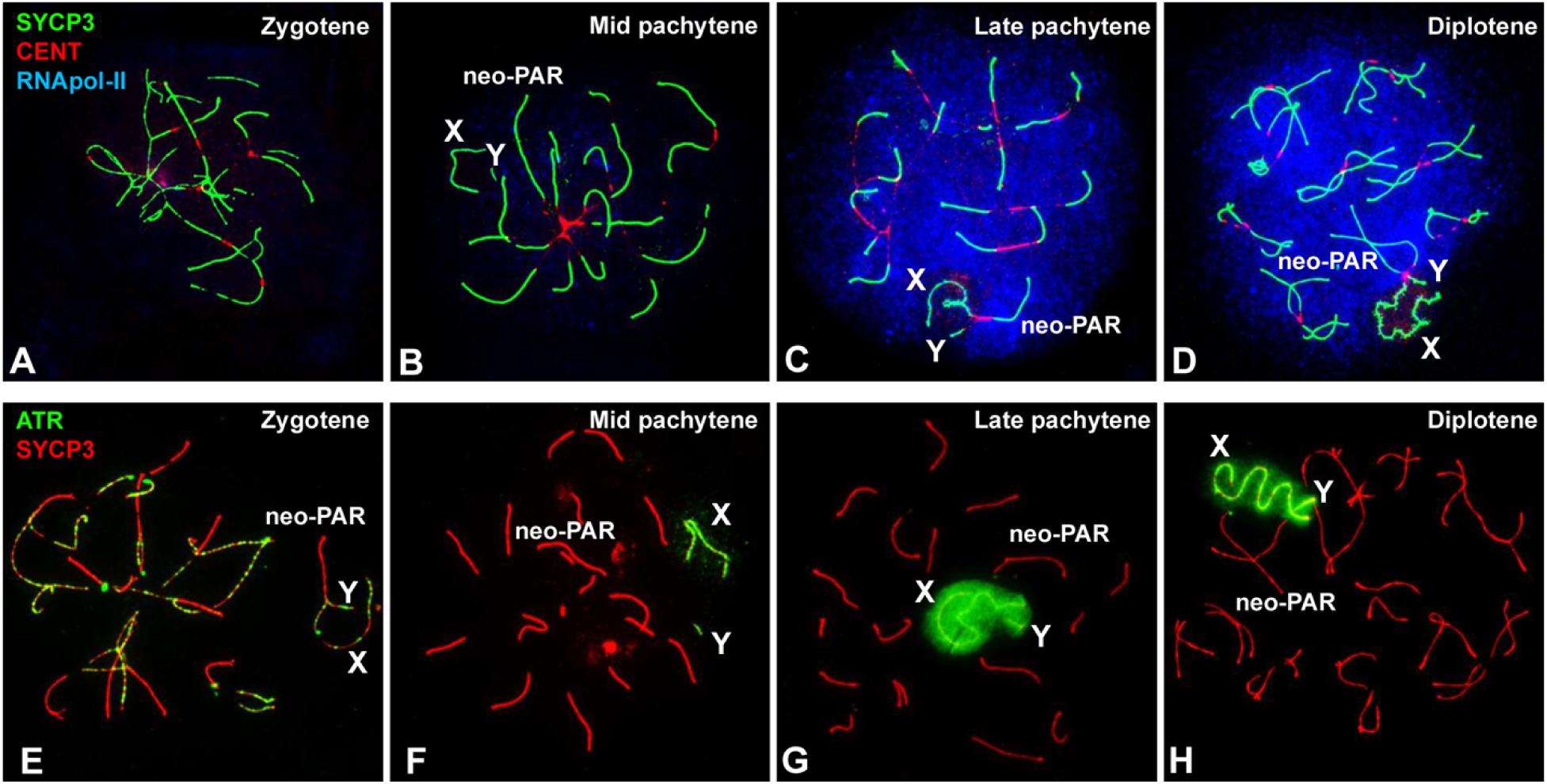
Inactivation of sex chromosomes. **A-D**. Spread spermatocytes at different stages of prophase-I labelled with antibodies against SYCP3 (green), centromeres (red) and RNA polymerase-II (blue). Non-homologous segments of the sex chromosomes (X, Y) and the neo-PAR are indicated. **A.** RNA pol-II is absent from the nucleus during zygotene. **B.** At mid pachytene, a weak RNA pol-II signal is detected in several areas of the nucleus. **C.** By late pachytene, the nuclear RNA pol-II signal has greatly increased and is maintained through diplotene (**D**). In contrast to the neo-PAR and autosomes, the non-homologous segments of the sex chromosomes (X, Y) are mostly or completely devoid of RNA pol-II. **E-H**. Spread spermatocytes at different stages of prophase-I labelled with antibodies against SYCP3 (red) and ATR (green). **E.** At zygotene, ATR foci are observed along the unsynapsed regions of the autosomes and sex chromosomes. **F**. At mid pachytene, ATR is only seen on the unsynapsed regions of the sex chromosomes. At late pachytene (**G**) and diplotene (**H**), the ATR signal has extended to the chromatin of the non-homologous segments of the sex chromosomes. The neo-PAR is not labelled by ATR.

### Recombination in the neo-PAR

We next assessed recombination in the neo-PAR during male meiosis by analyzing the frequency and distribution of chiasmata using the cytological marker MLH1 [58] to identify the location of crossovers along chromosomes (supplementary Fig. S5). We divided the neo-PAR into ten equivalent intervals and the frequency of MLH1 foci in each interval was recorded in a total of 87 spermatocytes from three *M. minutoides* males. We found that the neo-PAR systematically displays one or two chiasmata (mean=1.02), but that they are not distributed all along the chromosome arm (Fig. 4A). Notably, MLH1 is not observed on interval 1 to 3, and intervals 4 and 5 have a relatively low recombination frequency. Thus, MLH1 foci are strictly excluded from the same segments that show delayed synapsis and DNA repair. To discard the possibility that chiasmata displacement were a general effect of Rb translocations in *M. minutoides*, we additionally analyzed the distribution of MLH1 in the longest autosomal arm. The chiasmata frequency for this arm was slightly higher (mean=1.07 in 91 spermatocytes) and no exclusion was found in any of the chromosomal segments (Fig. 4A), which suggests that the Rb translocation alone is not responsible for the lack of recombination in the proximal region of the neo-PAR. Finally, to discard the possibility that the absence of chiasmata in this region is due to an intrinsic feature of the autosome translocated with the sex chromosomes in *M. minutoides*, we also evaluated the distribution of MLH1 in chromosome 1 in the closely related pygmy mouse *Mus mattheyi.* In this species sex chromosomes are not fused to chromosome pair 1. Therefore, it presents the ancestral sex chromosome configuration found in most pygmy mice, i.e., they are achiasmate, while chromosome 1 remains autosomal (supplementary Fig. S5). We recorded MLH1 foci along this chromosome in a total of 70 spermatocytes from three males and found a slight increase in number of chiasmata (mean=1.10) compared with the neo-PAR of *M. minutoides*. More relevantly, in *M. mattheyi*, MLH1 is distributed all along chromosome 1, except in interval 1, which is close to the centromere, and interval 3 and 4 have the highest recombination frequencies (supplementary Fig. S5).

**Figure 4.**
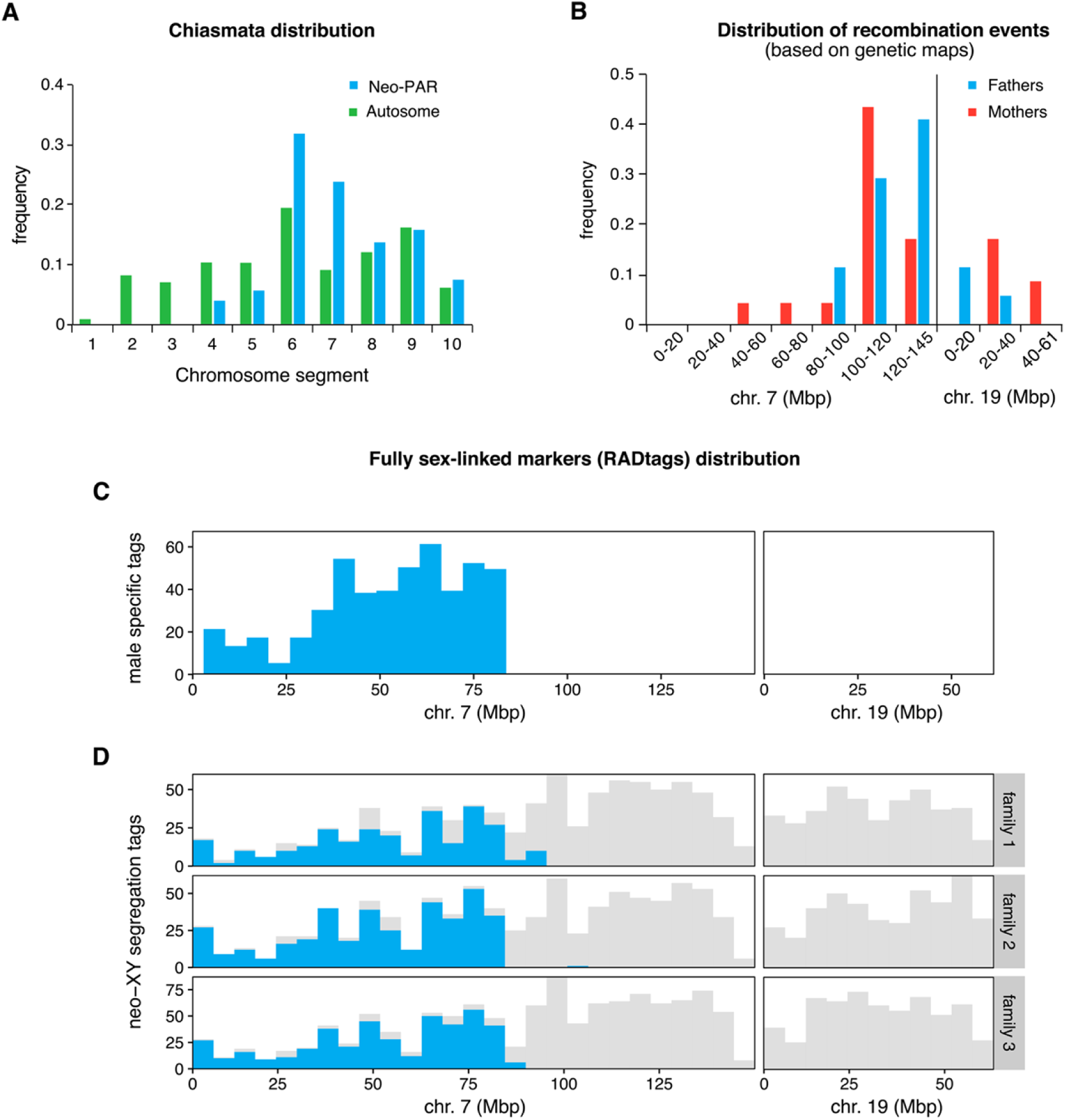
Recombination and position of fully sex-linked genetic markers on the neo-sex chromosomes. **A**. Distribution of MLH1 foci along the neo-PAR in *M. minutoides* (blue) and the longest autosomal arm (green). The chromosomes have been divided into 10 equal segments from the centromere (1) to the distal telomere (10). **B.** Distribution of recombination events across the neo-sex chromosomes in the parents of families 1 and 3 (red: mothers, blue: fathers) based on the recombination maps build independently for the four parents. **C.** Distribution of male-specific RADtags across the neo-sex chromosomes, according to their mapping position to *M. musculus domesticus* chromosomes 7 and 19. **D.** Distribution of tags carrying SNPs with a fully sex-linked segregation pattern along the neo-sex chromosomes, according to their mapping position to *M. musculus domesticus* chromosomes 7 and 19 (shown in blue). Tags that carry at least one SNP, regardless of their transmission pattern, are shown in grey.

These observations suggest that the proximal third of the Y neo-PAR in *M. minutoides* no longer recombines and, therefore, has become fully sex-linked. To confirm this result, we used restriction site-associated DNA sequencing (RAD-seq), a technique commonly used to detect non-recombining regions on sex chromosomes through the identification of sex-linked genetic markers [59-62]. Three full-sibling families (hereafter called 1, 2 and 3) were sequenced (total: 65 individuals, including parents and sexed offspring). The final number of high-quality genetic markers (single nucleotide polymorphisms, hereafter SNPs) retained for families 1, 2 and 3, respectively, was 29,518; 30,202 and 44,131 across 22,777; 24,131 and 34,426 RADtags. This dataset was screened for fully sex-linked tags and markers using two approaches derived from those used by Brelsford et al. [63]. The first method identifies sex-limited RADtags specific to the Y chromosome, i.e., sequences found in all males but absent in all females. The second method screens for fully sex-linked SNPs present in both sexes that follow a male-heterogametic segregation pattern (heterozygous father and sons, homozygous mother and daughters). The results of the two approaches are displayed in Table 1 and Fig. 4B-D. Across all families, we found over a thousand RADtags that were present only in males, and several hundred markers that follow a sex-linked segregation pattern in each family. All candidate RADtags with full sex linkage were aligned to the *M. musculus domesticus* reference genome to determine if the neo-PAR in *M. minutoides* is enriched in male-limited alleles (as expected if recombination was suppressed). The proximal region of the neo-PAR in *M. minutoides* is homologous to chromosome 7 in *M. musculus domesticus* and its distal region to chromosome 19 [64]. Therefore, we focused on tags mapping to these two chromosomes of the reference genome. Of the candidate RADtags that were successfully aligned, over 90% mapped to chromosome 7, while none mapped to chromosome 19 (Table 1, see supplementary Table S1 for the full record of alignment hits). Candidate RADtags were not distributed evenly along chromosome 7: they all aligned to the proximal region of the chromosome (Fig. 4C and D). Among the three families, the most distal fully sex-linked markers that aligned to chromosome 7 were consistently found in a similar location (between 83.5Mb and 92.7Mb, Table 1). RADtag density in this region, which also contained non-sexed-linked SNPs, was high (Fig. 4C and D). This result suggests that we are accurately describing the position of the most proximal recombination events on the neo-PAR during male meiosis. Assuming no major intra-chromosomal rearrangements between the *M. minutoides* neo-PAR and *M. musculus domesticus* chromosome 7, these results suggest that up to 83.5 Mbp of the neo-PAR is a non-recombining region. Interestingly, we also found a striking sex difference in the proportion of heterozygous sites along this region (supplementary Fig. S6). In males, the proportion of heterozygous sites along this region is similar to that found across the distal part of the neo-PAR in both sexes; however, in females, it is close to 0. This pattern conforms to the expectations for an X chromosome-specific region: as a consequence of the absence of recombination with the Y chromosome in males, effective population size decreases, and polymorphism loss driven by genetic drift is more likely, resulting in reduced heterozygosity.

**Table 1.**
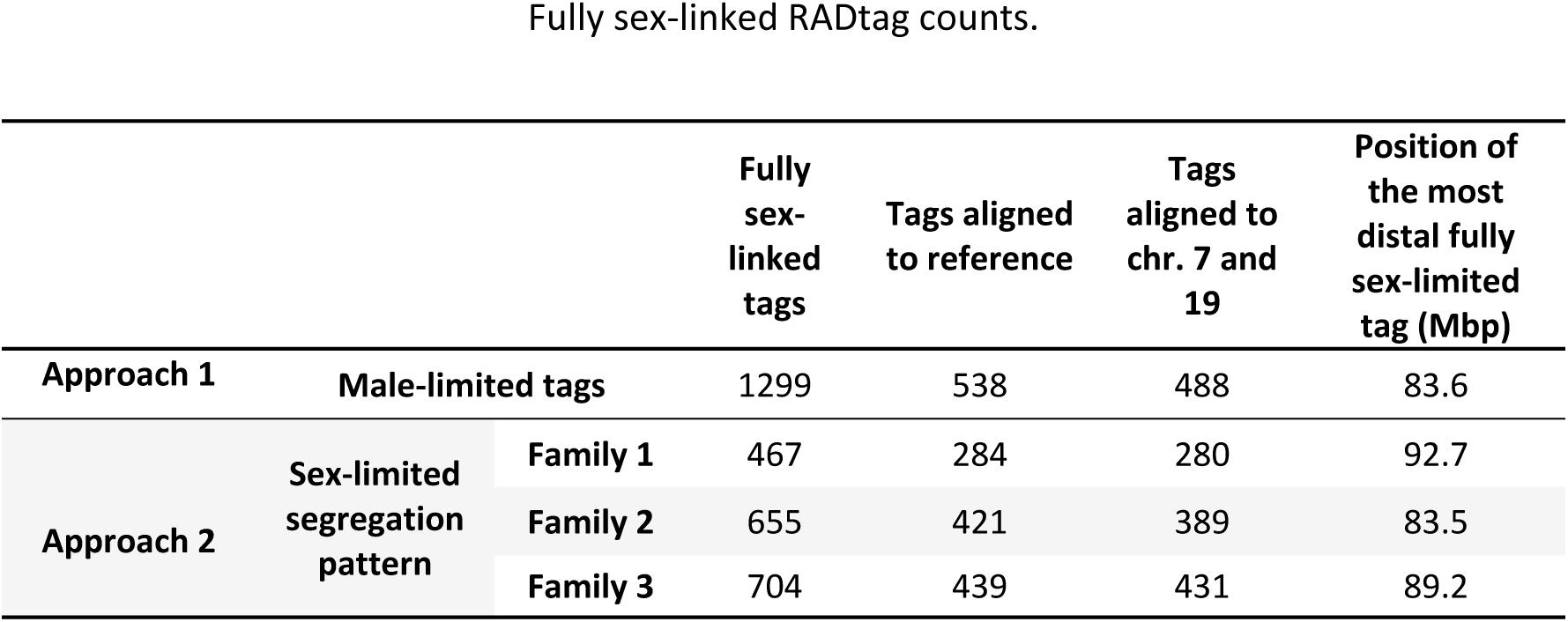
Fully sex-linked RADtag counts. The positions of the most distal fully sex-linked tags are based on their map position relative to chromosome 7 of the *M. musculus domesticus* reference genome (Mbp: megabase pairs).

For families 1 and 3, all of the genetic markers found on RADtags aligning to reference chromosomes 7 and 19 were used to build a recombination map of the neo-PAR for each parent in order to infer the distribution of recombination events based on the genetic data (unfortunately, family 2 had too few offspring to successfully build recombination maps). The segregation data reveal a clear sex difference in crossover patterns (Fig. 4B, supplementary Fig. S7 and supplementary Table S2). In fathers, crossovers occurred only in the distal region of the neo-PAR, and all markers that aligned to the proximal region of chromosome 7 consistently segregated together (i.e., they are at the same position on the recombination map). This result is in line with chiasmata distribution observed during male meiosis (Fig. 4A). In contrast, in females, several recombination events were observed within the proximal region (supplementary Table S2), alike on male autosomes and chromosome 1 of *M. mattheyi* (Fig. 4A and supplementary Fig. S5). It is worth noting that the recombination rate of this region is likely underestimated in females. Indeed, only heterozygous markers are useful to build recombination maps, therefore, crossovers were potentially missed due to the extremely low heterozygosity found in this region.

### Sex chromosomes display a dual mechanism of association at metaphase-I

Finally, we analyzed sex chromosome segregation during the first meiotic division by following the distribution of SYCP3 and γH2AX in spermatocyte squashes. SYCP3, which was present along chromosomes during prophase-I (see Fig. 2), relocalizes to the region of contact between sister chromatids at metaphase-I and accumulates at the centromeres (Fig. 5A-D). The pattern of SYCP3 localization allows the precise identification of bivalent orientation. As expected, we observed that the X and Y chromosomes are linked by at least one chiasma in the neo-PAR. The non-homologous segments, which are identifiable by the intense γH2AX labelling, never show a chiasma. However, we found they can display different configurations at metaphase-I: 1) they can appear associated, forming a single chromatin body labelled with γH2AX (Fig. 5A); 2) alternatively, they can appear as separate chromatin masses that are connected to each other by either SYCP3- or γH2AX-positive filaments or both (Fig. 5B-C) or 3) they can appear as two distinct masses that are not in contact, such that the only association between the sex chromosomes is through the chiasma in the neo-PAR (Fig. 5D). The proportion of the different configurations displayed is not equal. Of the metaphase cells analyzed (n=40 in two individuals), 75% have some type of connection between the non-homologous segments of X and Y, while only 25% are completely separated.

**Figure 5.**
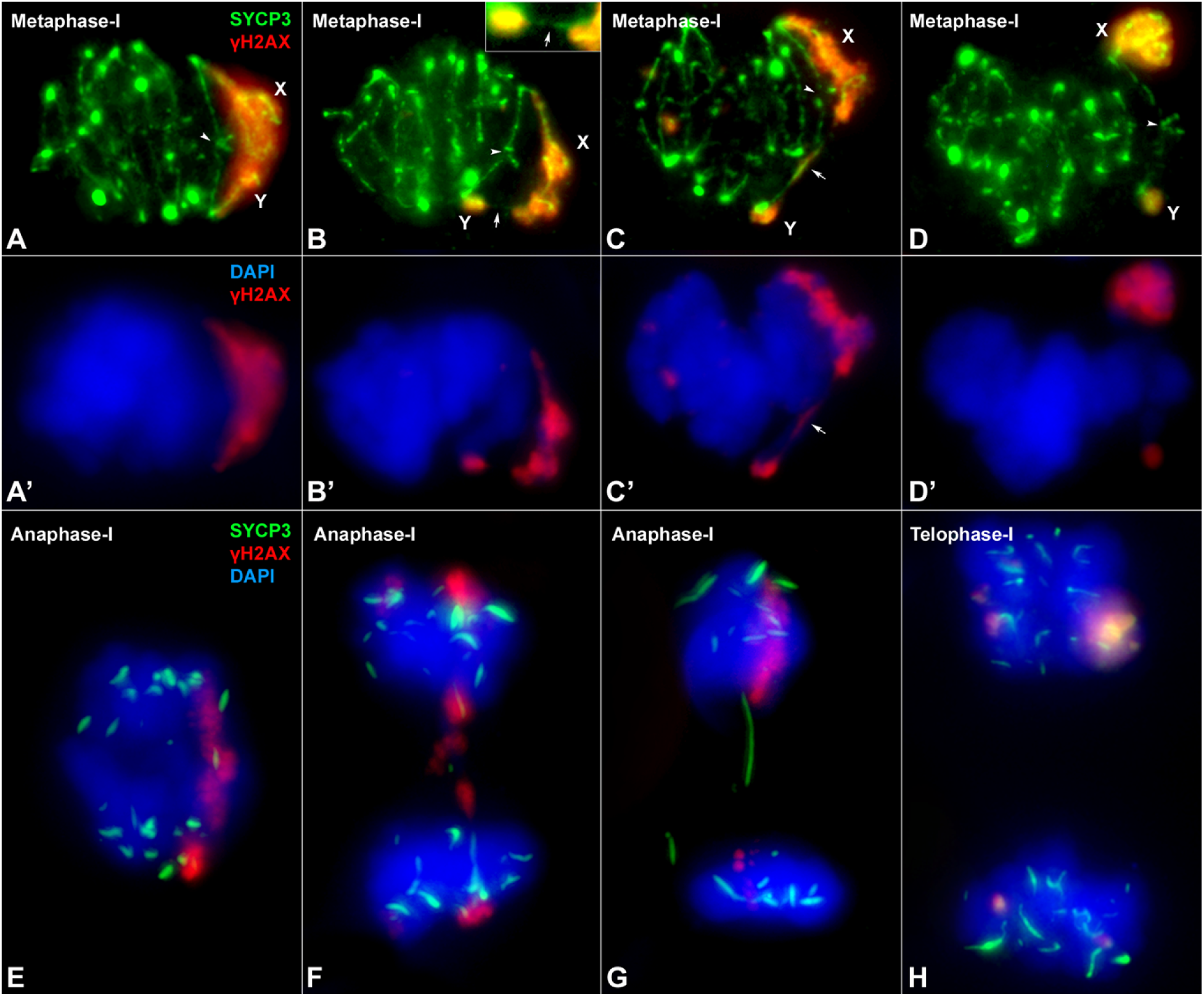
Sex chromosome segregation. Squashed spermatocytes labelled with antibodies against SYCP3 (green) and γH2AX (red) and counterstained with DAPI (blue). Non-homologous segments of the sex chromosomes (X, Y) are indicated. **A-D’**. Metaphase-I. Different configurations of the sex bivalent can be observed. The neo-PAR always shows a chiasma (arrowheads), however, the non-homologous segments can associated with each other by forming a common chromatin mass (**A-A’**), or through a SYCP3-positive filament (arrow) (see enlarged detail on the top) (**B-B’**) or a γH2AX-positive filament (arrow) (**C-C’**). Alternatively, these segments can appear completely separated (**D-D’**). **E-G**. Anaphase-I. γH2AX connections (E) and/or SYCP3 (F, G) filaments are observed between segregating chromosomes during early anaphase, resembling the associations observed at metaphase-I. At telophase-I (**H**), no lagging or mis-segregated chromosomes are observed.

Once all the chromosomes have aligned along the cellular equator, they begin to migrate at the onset of anaphase-I. At this stage, we also observed different configurations in relation to sex chromosomes segregation. In some anaphase-I cells, the two sex chromosomes are connected by γH2AX-positive filaments (Fig. 5E), while in others, they are connected by SYCP3 filaments that also appear to be associated with chromatin regions still marked with γH2AX (Fig. 5F) or by filaments that cross from one pole to the other (Fig. 5G). These configurations seem to correspond to the remnants of the ones observed at metaphase-I. In any case, at telophase-I, no chromosomes are lagging (Fig. 5H), indicating that the sex chromosomes segregate correctly.

## DISCUSSION

The study of the early stages of sex chromosome evolution in mammals has been extremely elusive, mainly because, the mammalian sex chromosomes are at a very late stage of differentiation. Therefore, this system cannot provide information on the initial steps of recombination cessation or its meiotic causes and consequences. However, autosomes translocated to sex chromosomes, as a part of an addition-attrition cycle [8], represent a unique model to investigate early sex chromosome differentiation and evolution.

### The neo-PAR in *M. minutoides* already displays some, but not all, meiotic sex chromosome features

Delayed synapsis and DNA damage repair in the neo-PAR of *M. minutoides* indicate the “sexualization” of this chromosome segment, as these are meiotic features typical of differentiated sex chromosomes [33]. Structural constrains may influence these processes. The length of the neo-PAR, the extreme size difference of the non-homologous segments of the X and Y chromosomes and the fact that all chromosome ends remain attached to the nuclear envelope force the X chromosome to bend significantly (see Fig. 2F), which may be the cause of the synapsis delay, as has been reported for other chromosomal rearrangements [65, 66]. This delay may be responsible for the accumulation of unresolved DNA damage intermediates on the unsynapsed regions of the neo-PAR. Conversely, delayed synapsis might be caused by deficient homology recognition between the chromosomes in the proximal region of the neo-PAR. In any case, these findings reveal that delayed synapsis and DNA damage resolution can appear prior to any morphological differentiation and, thus, represent the first processes affected during the initial steps of sex chromosome divergence.

However, other meiotic features might be more difficult to achieve. Most relevantly, the neo-PAR in *M. minutoides* is not subjected to MSCI, similar to the behavior found in other species with sex chromosomes to autosomes translocations [19, 20, 46, 67]. It has been argued that the interposition of heterochromatic blocks or other structural barriers might prevent the extension of inactivation to regions of autosomal origin [19, 20]. Although we cannot rule out the possibility that extended centromeres prevent MSCI expansion in *M. minutoides*, it is widely accepted that MSCI is mainly regulated by the presence of unsynapsed regions [68, 69]. Thus, we propose that the neo-PAR in *M. minutoides* is not subjected to MSCI because it is able to complete synapsis just before or right as the transcription burst occurs during pachytene. Otherwise, meiotic progression would be seriously compromised as the extension of inactivation may affect genes in the neo-PAR crucial for meiosis [43, 45]. In most mammals, MSCI is possible because, during evolution, their sex chromosomes have been emptied of meiotic-crucial genes and because many X-linked genes have autosomal paralogs originated by retrotransposition and, expressed only during meiosis [70-72]. However, it is likely that the translocation event in *M. minutoides* is too recent to have reached this step in the evolution of sex chromosomes. It is possible that the neo-PAR still carries meiotic-crucial genes and, in which case MSCI would be greatly deleterious. A full assembly and annotation of the *M. minutoides* genome would shed light on the presence of such genes in the neo-PAR.

### Recombination is suppressed in the proximal region of the neo-PAR

Cessation of recombination is the cornerstone of sex chromosome evolution. It triggers the genetic differentiation of the X and Y chromosomes [3, 5], mediated in particular by the degeneration of Y due to genetic isolation [2, 73]. Based on our combined study of chiasmata distribution in spermatocytes and segregation of genetic markers, we found no evidence of recombination between X and Y over a large region encompassing the proximal third of the neo-PAR. MLH1-independent recombination is possible (although rare) [74], therefore, we cannot firmly conclude that recombination is completely halted in this region during male meiosis. However, our results do suggest that recombination is, at least, very limited. Furthermore, the presence of thousands of fully sex-linked genetic markers (particularly, RADtags found only in males) in this region suggests that the Y neo-PAR carries alleles not found on the X. This observation, along with the highly reduced level of polymorphism observed in the exact same region of the X chromosome (see supplementary Fig. S7), conform to the early stages of X-Y differentiation and support the complete recombination suppression in the proximal segment of the neo-PAR. However, theoretical studies show that even rare events of recombination are enough to prevent genetic differentiation of sex chromosomes [75]. Another interesting and rare feature of *M. minutoides* is the presence of XY females: XY females carry a feminizing X that differs from the X found in males [76]. This feminizing X is also fused to chromosome 1 in the majority of *M. minutoides* populations and may show recombination in the neo-PAR. However, Baudat et al. [77] recently investigated *M. minutoides* meiosis in XY females and they reported that the proximal segment of the neo-PAR of the Y chromosome is always involved in heterologous synapsis (with no MLH1 foci). Consistent with our RAD-seq analyses, their results strongly suggest that recombination is also highly reduced or suppressed in the proximal segment of the neo-PAR in XY females. Screening for sex-linked markers in males and females from natural populations would reveal the extent of sex linkage in this region of the neo-PAR, potentially confirming our findings that it is fully sex-linked.

Although recombination suppression between sex chromosomes is common in many animal groups, the factors involved in the process are still unclear. Chromosomal rearrangements have often been invoked as possible proximal mechanisms, and sexually antagonistic selection as a potential ultimate factor [2-5, 73], but these mechanisms are still largely debated [78, 79]. In fact, it has been shown that, in the Okinawa spiny rat *Tokudaia muenninki*, suppression of recombination can occur in the absence of large chromosomal rearrangements [41]. Although we cannot completely rule out that a large inversion on the Y neo-PAR blocks recombination in *M. minutoides*, we do not favor this possibility as the X and the Y neo-PAR display no differences in banding patterns [48]. Furthermore, meiotic effects of an inversion, such as the formation of synaptic loops [80], were not detected. We also doubt that sexually antagonistic selection is involved. This selective force could favor recombination suppression along the neo-PAR through the linkage of male-beneficial but female-detrimental alleles to the sex-determining region of the Y chromosome, even if they are highly detrimental to females. This is because linkage disequilibrium would make these alleles limited to males [81]. However, if the neo-PAR carried genes with male-beneficial/female-detrimental alleles, the existence of XY females would select against it. Furthermore, if such alleles were already fixed on the proximal region of the neo-Y, XY females would have decreased reproductive success, which is not the case [82].

Given the arguments above and the results presented here, we propose that additional structural factors likely play an important role in recombination suppression in *M. minutoides*. A first possible cause is the Rb fusion itself. Such events are known to reduce genetic exchanges close to the centromere, as repeatedly reported in *M. musculus domesticus* Rb models [50, 83]. This fits with the observation that the non-recombining region is the proximal part of the neo-PAR. However, recombination events were detected in the proximal region of Rb autosomes and in the neo-sex chromosomes in females, providing evidence that restricted recombination in males cannot be due solely to the Rb fusion. Second, synapsis delay could contribute to a perturbation in recombination timing, eventually leading to its suppression. A vast literature has reported that chromosomes, especially sex chromosomes, fail to recombine when synapsis or DNA repair are disturbed or delayed [27, 57, 84-87]. This synaptic impairment could drive the loss of recombination in regions distal to the centromere. If recombination suppression leads to divergence of the affected regions, with time, those regions would not be able to serve as a proper molecular template for DNA repair with the cognate chromosome. Concurrently, synapsis would be inefficient, creating a feedback loop between impairment of synapsis and recombination. Given this context, we propose that synapsis in the proximal region of the neo-PAR observed at late pachytene in *M. minutoides* may already be heterologous (Fig. 6) and, therefore, this region would be no longer be considered part of the neo-PAR. In line with this proposal, the proximal region of the Y neo-PAR usually engages in heterologous synapsis with other chromosomes during XY female meiosis [77]. The fact that the neo-PAR is still able to complete synapsis in male meiosis could be due to a process known as synaptic adjustment that is typical in the non-homologous regions of the X and Y chromosomes in most mammals [15, 32, 33, 80]. Importantly, heterologous synapsis does not promote reciprocal recombination, and DNA damage is usually repaired with the sister chromatid [88].

**Figure 6.**
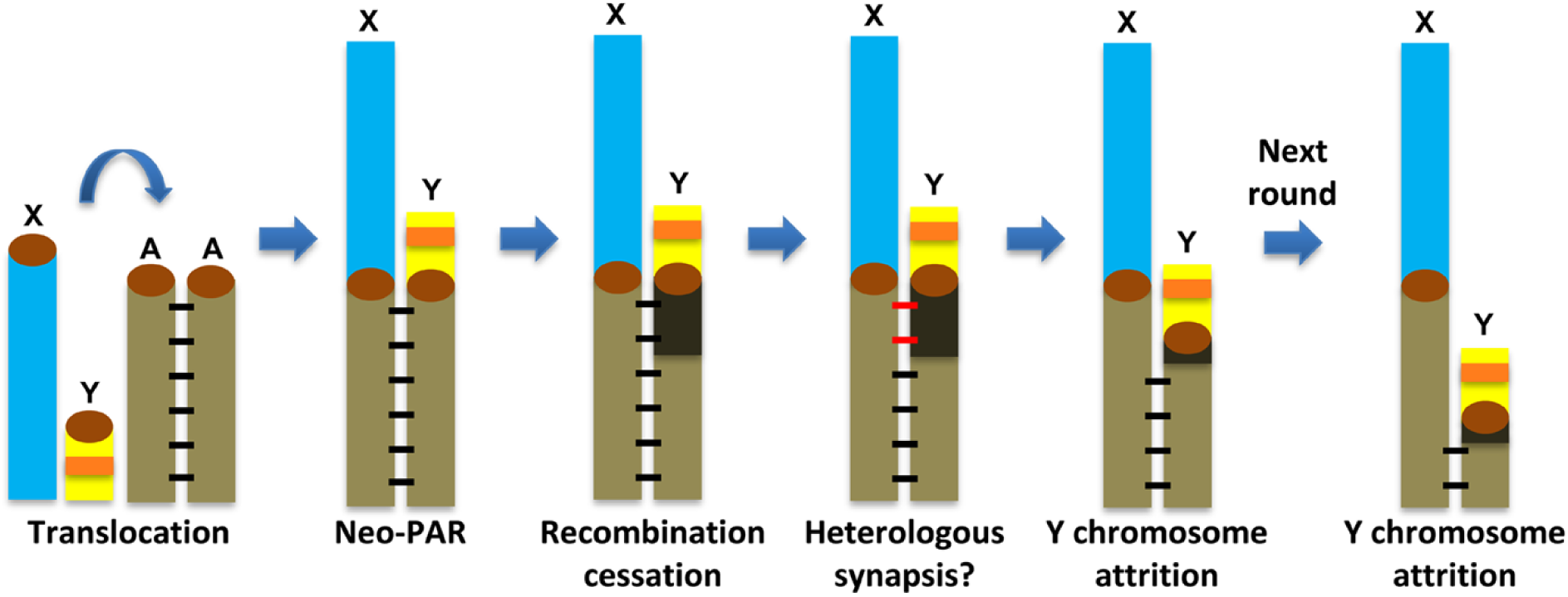
Schematic representation of a likely scenario for the evolution of the sex chromosomes in *M. minutoides* in the addition-attrition model. After the translocation event, a large neo-PAR region was formed, which shows complete synapsis (horizontal black bars) and recombination. This is followed by the first signs of modification: cessation of recombination and delay of synapsis and DNA repair, in the proximal segment of the neo-PAR. These processes could consequently lead to genetic differentiation of the Y neo-PAR (shadowed in dark brown) and heterologous synapsis of the differentiated region (red lines). A subsequent event of chromosome deletion could then lead to attrition and morphological differentiation of the Y chromosome. At this point, a new round of synapsis and recombination impairment could start over, promoting the recurrent attrition of the Y chromosome.

Our current knowledge of neo-sex chromosome evolution and differentiation is primarily due to extensive work on *Drosophila* (e.g. [89-92]. In this model, males do not recombine (male achiasmy), therefore recombination is immediately arrested after fusion to a Y chromosome. In contrast, the autosomes fused to sex chromosomes in mammals usually keep recombining in most, if not all, their full length. The only notable exception is in the black muntjac, where a large inversion on an X-autosome fusion led to recombination suppression and subsequent neo-Y degeneration [42]. Our findings in *M. minutoides* are thus significant since they represent one of the very rare cases in mammals, and the first that combine cellular and genetic approaches to highlight the early stages of neo-sex chromosome differentiation.

### A dual mechanism of sex chromosome association at metaphase-I

Another striking feature found in *M. minutoides* meiosis is the coexistence of a chiasma in the neo-PAR and an achiasmate association in the non-homologous segments. This feature, to our knowledge, has not been previously reported. The achiasmate association is most likely a relic of the one that the primitive sex chromosomes presented before the Rb translocation with chromosome 1 [16]. Indeed, it is analogous to mechanisms found in other mammalian species with achiasmate sex chromosomes, which typically involve the SYCP3 protein [11, 15, 93-95], or telomeric or heterochromatin-mediated associations [96-98]. The preservation of this mechanism, in addition to the chiasma in the neo-PAR, may merely be a side effect of the chromosomal translocation; on the other hand, it may prove to be evolutionary relevant. First, it may contribute to a more faithful segregation of the giant sex chromosomes during anaphase-I. Second, it may allow sex chromosomes to segregate properly under a scenario of chromosomal changes. *Mus minutoides* is extraordinarily variable in terms of chromosome number owing to different Rb rearrangements [48, 51] and populations in which only the Y chromosome has translocated with chromosome 1 have been found [49]. Phylogenetic analyses of these populations indicate that this sex chromosome configuration is a derived feature that originated by a fission of Rb (X.1), giving rise to a X1×2Y chromosome system in which X1 is the ancient X and X2 is the unfused chromosome 1. In the absence of a reliable mechanism to properly segregate the achiasmate X1 chromosome, one might expect the transmission of this chromosome to be compromised; however, the relic achiasmate association likely increased the probability of the fission being efficiently transmitted to these populations. An analogous preadaptative role of achiasmate modes of association has also been postulated in voles [15].

## CONCLUSIONS

The results presented here offer new clues to understand the evolution of sex chromosomes in mammals and they place meiosis as a pivotal factor. Structural constraints, such as differences in size, may condition the success of synapsis and recombination during meiosis. These elements mainly act after sex chromosomes have achieved a certain degree of morphological differentiation and, therefore, are likely more relevant during the addition-attrition cycle of sex chromosome evolution (Fig. 6). This could explain why sex chromosomes in eutherian mammals, which underwent initial differentiation [6], rapidly diverged even after the recurrent addition of autosomal chromosomes in the addition-attrition cycle [8]. On the other hand, without an initial stage of morphological differentiation, sex chromosomes could remain almost identical for very long periods, as is the case in many fishes, amphibians, reptiles and birds [2]. Meiotic studies of the sex chromosomes in these vertebrates could be useful to test this hypothesis.

Conversely, meiotic factors may also introduce restrictions on sex chromosome differentiation. First, attrition of the neo-Y causes a reduction of gene dose in males for the genes present in the segment involved, a process that could be deleterious for male meiosis. Second, sex chromosome differentiation leads to a process of MSCI, which potentially requires the translocation/transposition of many genes from the neo-PAR to the autosomes. This might be a difficult threshold to cross and could help to explain why old sex chromosomes in many vertebrates have barely achieved detectable morphological differentiation. We propose that the genetic and morphological constitution of each sex chromosome pair and the balance between meiotic factors promoting and hampering sex chromosome divergence result in an acceleration or delay in the evolution of these chromosomes.

## MATERIAL AND METHODS

### Animals

*Mus minutoides* and *M. mattheyi* were bred in captivity at the CECEMA facilities of Montpellier University. The colony was established from wild-caught animals as previously reported and maintained under standard conditions [77]. Males of both species were sacrificed by cervical dislocation and their testes processed for immunocytology. All experiments were conducted according to ethical rules established by the Institut des Sciences de l’Evolution of Montpellier and the Universidad Autónoma de Madrid (Ethics Committee Certificate CEI 55-999-A045).

### Immunofluorescence

For spreads and squashes, we followed the procedure described, respectively, by Peters et al. [99] and Page et al. [100]. Slides were incubated with primary antibodies diluted in PBS (137 mM NaCl, 2.7 mM KCl, 10.1 mM Na2HPO4, 1.7 mM KH2PO4, pH 7.4) overnight at room temperature in a moist chamber. The following primary antibodies and dilutions used were: mouse anti-SYCP3 (Abcam 97672), 1:100; rabbit anti-SYCP3 (Abcam 15093), 1:100, rabbit anti-histone H2AX phosphorylated at serine 139 (γH2AX) (Abcam 2893), 1:1000; mouse anti-γH2AX (Upstate 05-636), 1:1000; rabbit anti-RAD51 (Santa Cruz 8349), 1:50; mouse anti-MLH1 (Pharmingen 550838), 1:100; a human serum that recognizes centromere proteins (Antibodies Inc. 15-235), 1:100; mouse anti-RNA polymerase II phosphorylated at serine 2 (Abcam 24758), 1:100; and goat anti-ATR (Santa Cruz SC-1887), 1:100. After rinsing in PBS, the slides were incubated with the appropriate secondary antibodies diluted to 1:100 in PBS for one hour at room temperature: donkey anti-mouse, donkey anti rabbit, donkey anti-goat and goat anti-human, conjugated with Alexa 350, Alexa 488, Alexa 549 (Invitrogen), Cy3 or Dylight 649 (Jackson ImmunoResearch Laboratories). Slides were counterstained with DAPI, when needed, and mounted with Vectashield (Vector).

Slides were observed on an Olympus BX61 microscope equipped with a motorized plate in the Z axis. Images were obtained with an Olympus DP61 camera and processed with Adobe Photoshop 7.0 software. For the squashes, several optical sections were recorded for each cell. The stack files were processed with ImageJ to build 3-dimensional reconstructions as previously described [11, 40].

### Chiasmata distribution

Chiasmata distribution along chromosomes was analyzed in 91 spermatocytes from three *M. minutoides* males and 72 spermatocytes from three *M. mattheyi* males. The length of the neo-PAR and the largest autosomal arm (which according to the reported karyotype [48, 77] corresponds to chromosome Rb 2:10) in *M. minutoides*, as well as chromosome 1 in *M. mattheyi*, was measured using the free hand tool in ImageJ. These segments were then divided into 10 equally distant intervals from the centromere to the distal telomere. Then, we measured the distance between the centromere and the MLH1 foci, assigning each focus to its corresponding interval.

### Sample collection and RAD-seq library preparation

We generated double digest RAD-seq data for three *M. minutoides* full-sibling families bred in captivity (parents and offspring). We selected families according to the following three criteria: (i) XX sex chromosome complement of the mother, (ii) maximum number of sexed offspring and (iii) minimum relatedness between the two parents, which was assessed by a pedigree analysis of the breeding colony (see [82] for colony details). Families, referred to as family 1, 2 and 3 throughout the manuscript, were comprised of the following number of sons and daughters: 21 and 9 for family 1, 9 and 4 for family 2 and 12 and 3 for family 3 (the offspring in each family were born across multiple clutches). In family 2, two fathers sired offspring: the first sired 5 sons and 1 daughter, and the second 4 sons and 3 daughters. Tissue samples (muscle or tail) were collected from each individual and DNA was extracted using the Qiagen^®^ DNeasy^®^ Blood & Tissue Kit. The RAD-seq library was prepared following the protocol described by Brelsford et al. [59]. In brief, genomic DNA was digested with restriction enzymes EcoR1 and Mse1, adapters containing barcodes unique to each sample were ligated to the DNA fragments, which were then PCR amplified in 20 cycles. PCR products were pooled and approximately 350-450 bp fragments were isolated using gel-based size selection. The RAD-seq library was single-end (100 bp) sequenced on an Illumina HiSeq 2500.

### RAD-seq data processing and sample quality control

The quality of Illumina raw reads was checked using FASTQC v0.10.1 [101] and demultiplexed by individual barcode using the process_radtags module of STACKS v1.48 [102]. STACKS modules Ustacks (parameters –M 2, -m 3), Cstacks (-n 1) and Sstacks were run on the complete dataset. The module Populations, which filters RAD markers (SNPs) for varying presence/absence criteria across samples, was run for each family separately, first with relaxed parameters: markers were retained if present in at least half of the offspring with a depth of coverage ≥ 8. Under these parameters, we were able to identify any low-quality samples and/or potential family misassignments. In each family, over 100,000 RADtags were retained with an average tag coverage of approximately 20x. No low-quality samples were identified (based on estimations of per sample coverage, heterozygosity and tag absence); however, kinship analyses using the PLINK method of moments [103] revealed that two individuals had been incorrectly assigned to family 1 (kinship coefficients of 0.02 and 0.08 with the father and 0.17 and 0.16 with the mother). These two individuals were removed from further analyses. Because many markers with high coverage were present in the initial analyses, we used more stringent parameters for the final ones: markers were retained if present in both parents and all offspring with a depth of coverage ≥ 12. Additionally, we specified a minimum minor allele frequency of 0.05. The number of high-quality SNPs retained for families 1, 2 and 3, respectively, was 29,518; 30,202 and 44,131 across 22,777; 24,131 and 34,426 RADtags.

### Identification of fully sex-linked markers

The RAD-seq dataset was screened for fully sex-linked tags and markers using two approaches, both derived from those used by Brelsford et al. [63]. The first approach detects the presence of fully sex-linked tags: all RADtags were assessed for their presence or absence in both sexes and were only considered Y-limited if they were completely absent in females and present in at least 90% of males, across all individuals sequenced. We expect such loci to be found either on the non-homologous segment of the Y chromosome (ancestral Y), or in the neo-PAR, assuming it carries a fully sex-linked non-recombining region. Note that several mechanisms can result in such sex-specific loci: a deletion or a mutation in the restriction site on the X chromosome (leading to a sex-specific null allele), a mutation on Y that creates a novel Y-specific restriction site, or an under-merging of RADtags in Stacks due to a high level of divergence between X- and Y-limited alleles. We also screened for tags absent in males but present in at least 90% of all females. We found only four, suggesting a low level of false positives in our list of male-limited tags (consisting of 1299 tags). The second approach screens for fully sex-linked SNPs on tags present in both males and females, and it relies on the study of the segregation pattern of markers within full-sib families. Under this approach, a marker is considered fully sex-linked if it is (i) heterozygous in the father, (ii) homozygous in the mother, (iii) heterozygous in all sons and (iv) homozygous for the maternal allele in all daughters.

### Assessing the presence or absence of recombination along the neo-sex chromosomes

All RADtags were aligned to the *Mus musculus domesticus* GRCm38 reference genome using bwa-mem v0.7.12 with default parameters [104]. Alignments with a mapping quality lower than 30 were discarded. As the neo-sex chromosome in *M. minutoides* is homologous to chromosomes 7 and 19 of *M. musculus domesticus* [64], we evaluated whether there is an excess of fully sex-linked markers on these chromosomes. Additionally, we assessed the exact mapping position of these markers to determine if a specific region of the chromosome is enriched in sex-linked markers. The proportion of tags that could be successfully aligned with the reference genome differed by approach: approximately 40% for the male-limited tags, and 60% for the fully sex-linked tags present in both sexes. This discrepancy likely stems from the fact that many tags completely absent in females tend to be in the region of the ancestral Y chromosome that shares no homology with the ancestral X. This region is known to have a high rate of evolution (due to e.g. male mutation bias and rapid accumulation of repetitive DNA sequences) and, thus, likely shows higher sequence divergence between *M. minutoides* and *M. musculus domesticus* relative to other regions. We also evaluated the level of heterozygosity across all markers (regardless of sex linkage) along the neo-PAR in fathers and mothers. For each pair of parents, we identified all RADtags present in both individuals with a depth of coverage ≥ 8. Tags mapping to chromosomes 7 and 19 were screened for heterozygous sites in each parent, and the proportion of heterozygous sites was evaluated in 1-Mbp regions (based on the mapping positions of markers to the *M. musculus domesticus* chromosomes). Regions with reduced or no recombination should show a pattern of lower heterozygosity in females because of the reduced effective population size of the neo-X chromosome. In males, heterozygosity in non-recombining regions should be equivalent to or greater than that found along the neo-PAR. Finally, we took advantage of the full-sib family data and the availability of a reference genome to build parental recombination maps to identify genetic markers that segregate together and to assess crossover distribution in the two sexes. Specifically, using LepMap3 [105] with a minimum logarithm of odds (LOD) score of 2, and the Kosambi mapping function, we built sex-specific recombination maps for the neo-sex chromosomes based on all polymorphic markers mapping to *M. musculus* chromosomes 7 and 19 and evaluated the position of recombination events that occurred in each parent of the three families.

## ACKNOWLEDGEMENTS

We are grateful to Roberto Sermier for DNA extractions and help with the RADseq library preparation, Marie Challe for help in maintaining the breeding colonies, and Dr. C. García de la Vega for stimulating and fruitful discussions. This work was supported by from the Ministerio de Economía y Competitividad (Spain) (Grant number CGL2014-53106-P to JP), French National Research Agency (ANR grant SEXYMUS 10-JCJC-1605 and ANR program ANR-16-IDEX-0006 to FV), Del Duca Foundation from the Institut de France (“subvention scientifique”), and the Swiss National Science Foundation (grant 31003A 166323 to NP). A.G-F. was supported by a predoctoral fellowship from the Ministerio de Economía y Competitividad (Spain) and the European Social Fund (European Commission). Computations were performed at Vital-IT (www.vital-it.ch), a center for high-performance computing of the SIB Swiss Institute of Bioinformatics, and the Montpellier Bioinformatics Biodiversity (MBB) platform, supported by the LabEx CeMEB (ANR-IA-10-LABX-0004).

## SUPPLEMENTARY INFORMATION

**Figure S1.**
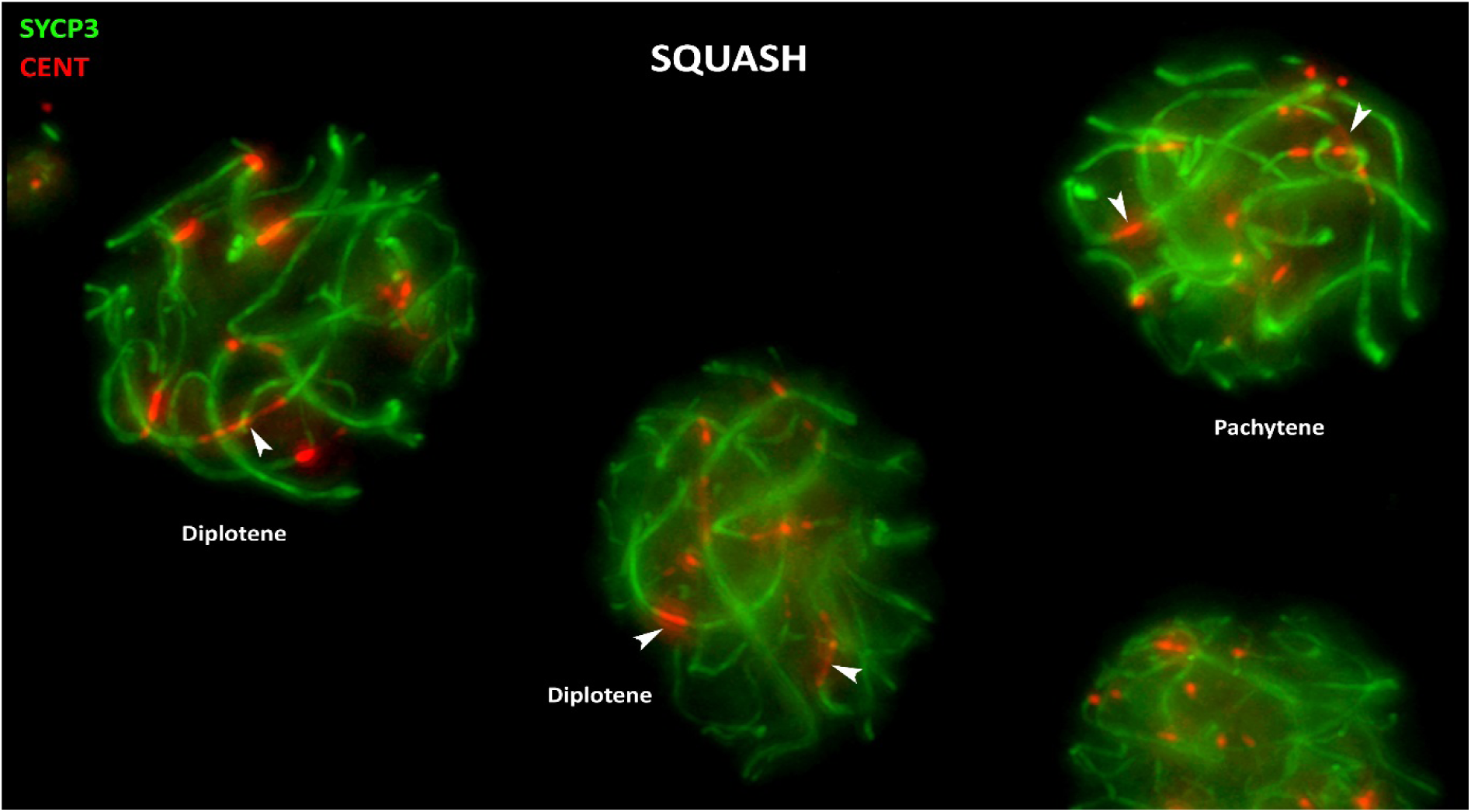
Squashed *M. minutoides* spermatocytes labelled with antibodies against SYCP3 (green) and centromeres (red). The stretching of centromeres (arrowheads) in pachytene and diplotene nuclei is observed clearly with this technique, which preserves the three-dimensional organization of the nucleus.

**Figure S2.**
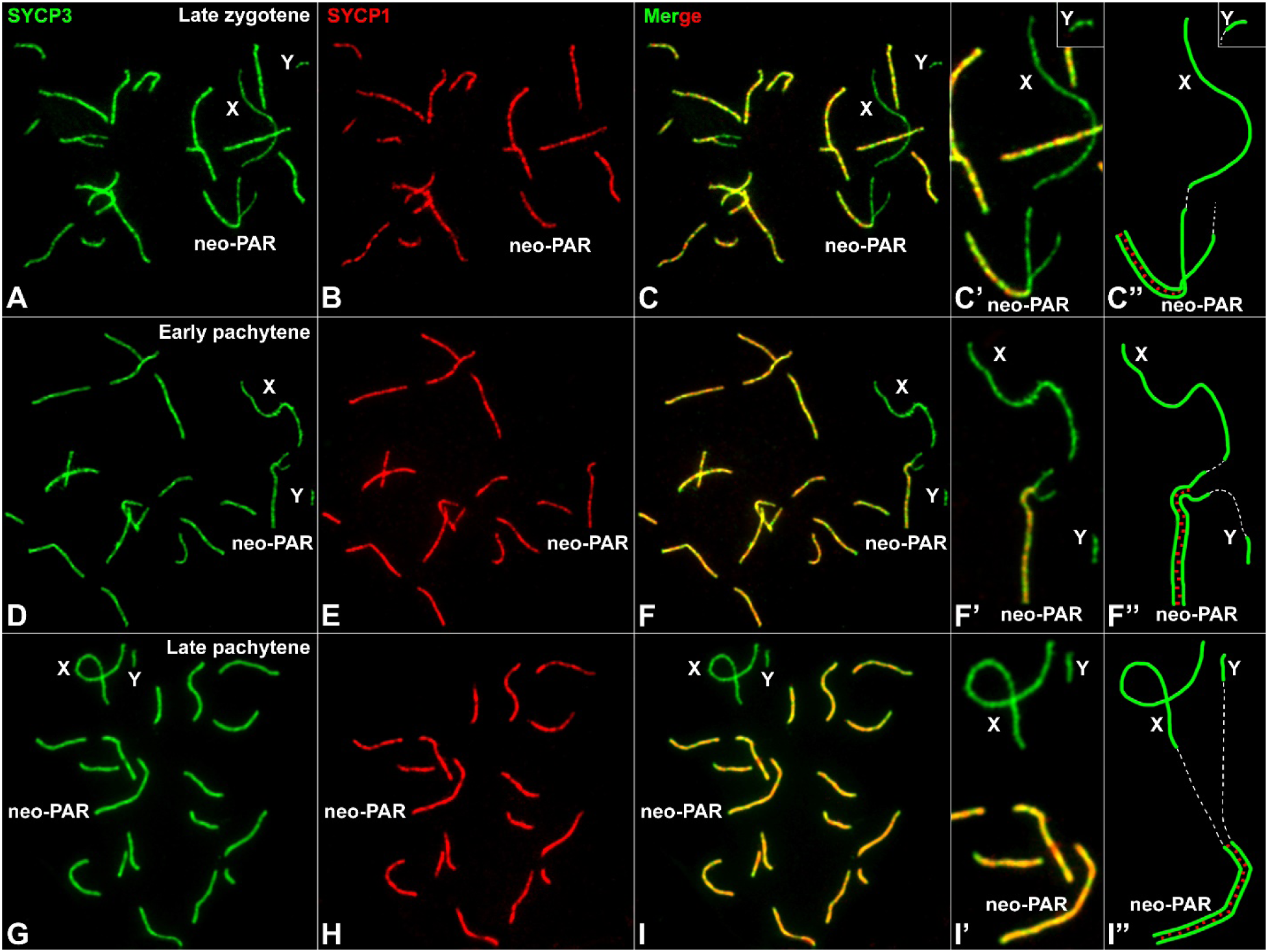
Synapsis in *M. minutoides*. Spread spermatocytes at different stages of prophase-I labelled with antibodies against SYCP3 (green) and SYCP1 (red). The heterologous segments of the sex chromosomes (X, Y) and the neo-PAR are indicated. **A-C’’**. Late zygotene. Most autosomes have completed synapsis. SYCP1 and SYPC3 colocalize along the synapsed regions (yellow in **C**), while only SYCP3 is detectable along unsynapsed ones. **C’**. An enlarged view of the sex chromosomes shown in C. **C’**’. Schematic representation of sex chromosome synapsis. LEs are represented in green, TF as short red lines and centromeres as white dashed lines. **D-F’’**. Early pachytene. Autosomes are fully synapsed but the neo-PAR of the sex chromosomes has not yet completed synapsis. **G-I’’**. Late pachytene. The neo-PAR is completely synapsed, but the non-homologous segments of the X and Y chromosomes remain unsynapsed and negative for SYCP1.

**Figure S3.**
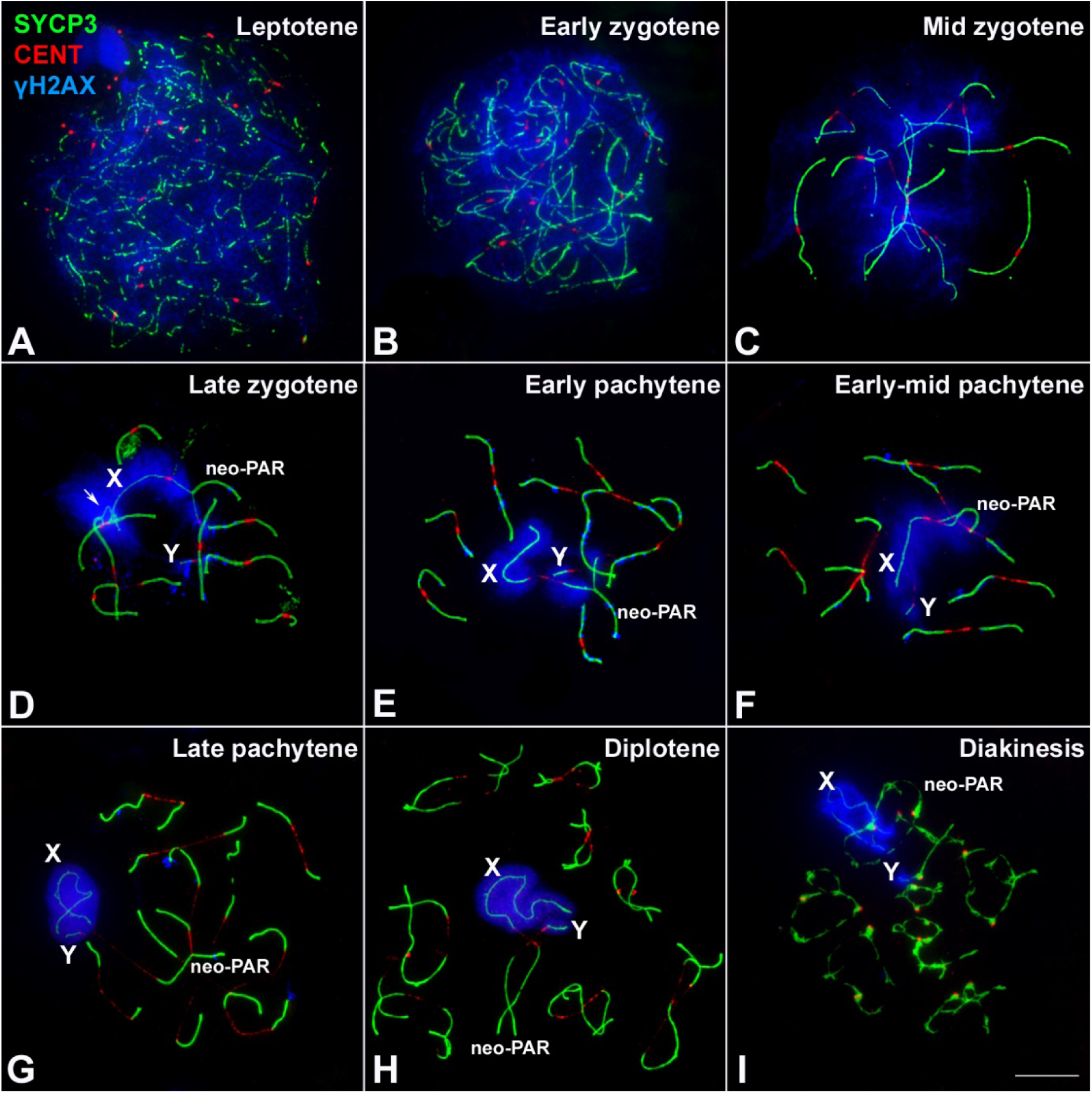
Complete sequence of synapsis and DNA repair in the sex chromosomes of *M. minutoides*. Spread spermatocytes at different stages of prophase-I labelled with antibodies against SYCP3 (green), centromeres (red) and γH2AX (blue). Sex chromosomes (X, Y) and the neo-PAR are indicated. **A**. Leptotene. SYPC3 forms short filaments, while γH2AX occupies the whole nucleus. **B-D**. Zygotene. SYCP3 forms thick filaments along the chromosomal regions that have completed synapsis. At early zygotene, the filaments are short, but elongate as synapsis proceeds. The γH2AX signal decreases during zygotene progression and, by late zygotene, only remains as a large focus on the unsynapsed regions of some autosomes and the sex chromosomes. Some small foci are also visible over synapsed autosomes. **E-G**. Pachytene. Autosomes have completed synapsis, but weak γH2AX signals can still be detected on them, mainly during early and mid pachytene. The neo-PAR of the sex chromosomes still remains partially unsynapsed at the transition from early to mid pachytene. During these stages, both the non-homologous segments and a portion of the neo-PAR are intensely labelled by γH2AX. By late pachytene, the neo-PAR appears completely synapsed. The non-homologous segments remain unsynapsed, though they associate with each other forming a sex body. **H**. Diplotene. The chromosomes initiate desynapsis and homologs remain associated at specific locations, likely chiasmata. γH2AX is observed only on the non-homologous segments of the sex chromosomes. **I**. Diakinesis. SYCP3 signal is irregular along the chromosomes and accumulates in some centromeric regions. γH2AX is only found on the non-homologous segments of the sex chromosomes.

**Figure S4.**
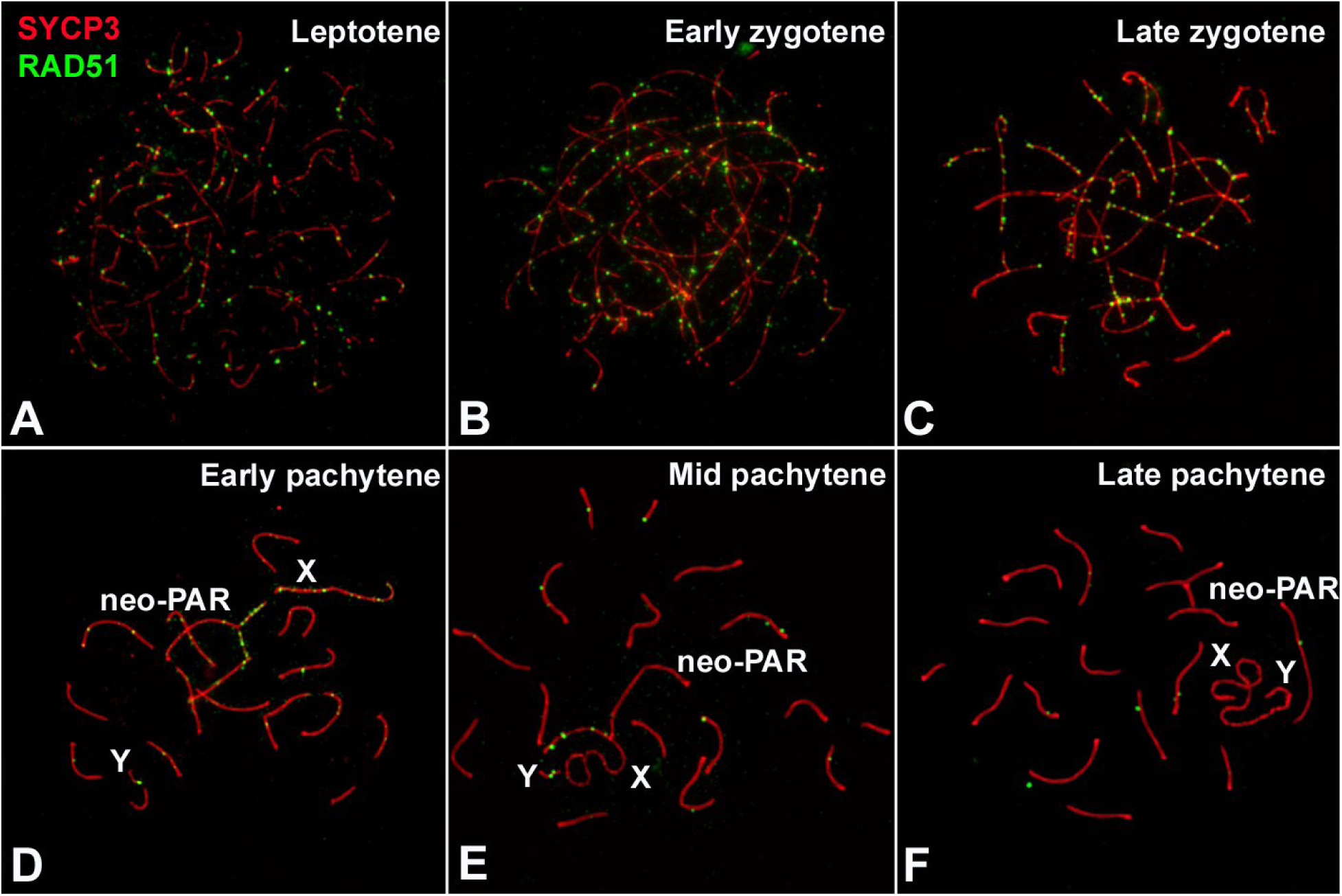
Complete sequence of DNA repair in the sex chromosomes of *M. minutoides*. Spread spermatocytes at different stages of prophase-I labelled with antibodies against SYCP3 (red) and RAD51 (green). Sex chromosomes (X, Y) and the neo-PAR are indicated.

**Figure S5.**
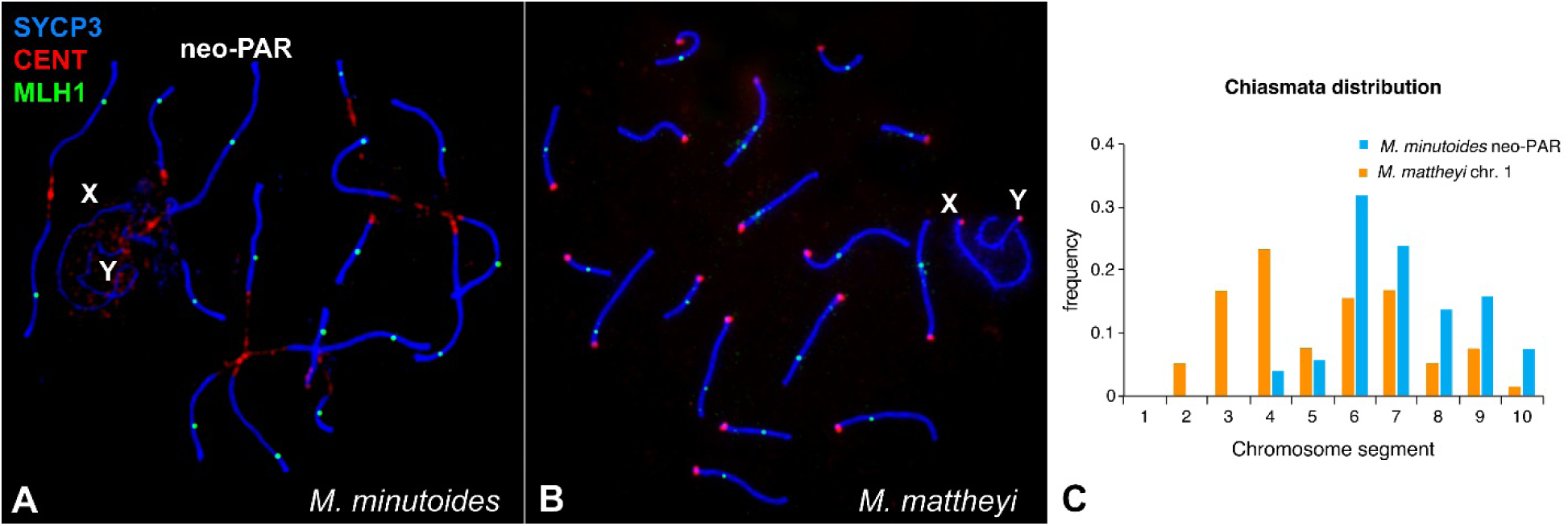
Spread spermatocytes of *M. minutoides* (**A**) and *M. mattheyi* (**B**) at pachytene labelled with antibodies against SYCP3 (blue), centromeres (red) and MLH1 (green). Sex chromosomes (X, Y) and the neo-PAR are indicated. Note the asynaptic behavior of sex chromosomes in *M. mattheyi*. **C**. Distribution of MLH1 foci along the neo-PAR in *M. minutoides* (blue) and the chromosome 1 in *M. mattheyi* (orange). The chromosomes have been divided into 10 equal segments from the centromere (1) to the distal telomere (10).

**Figure S6.**
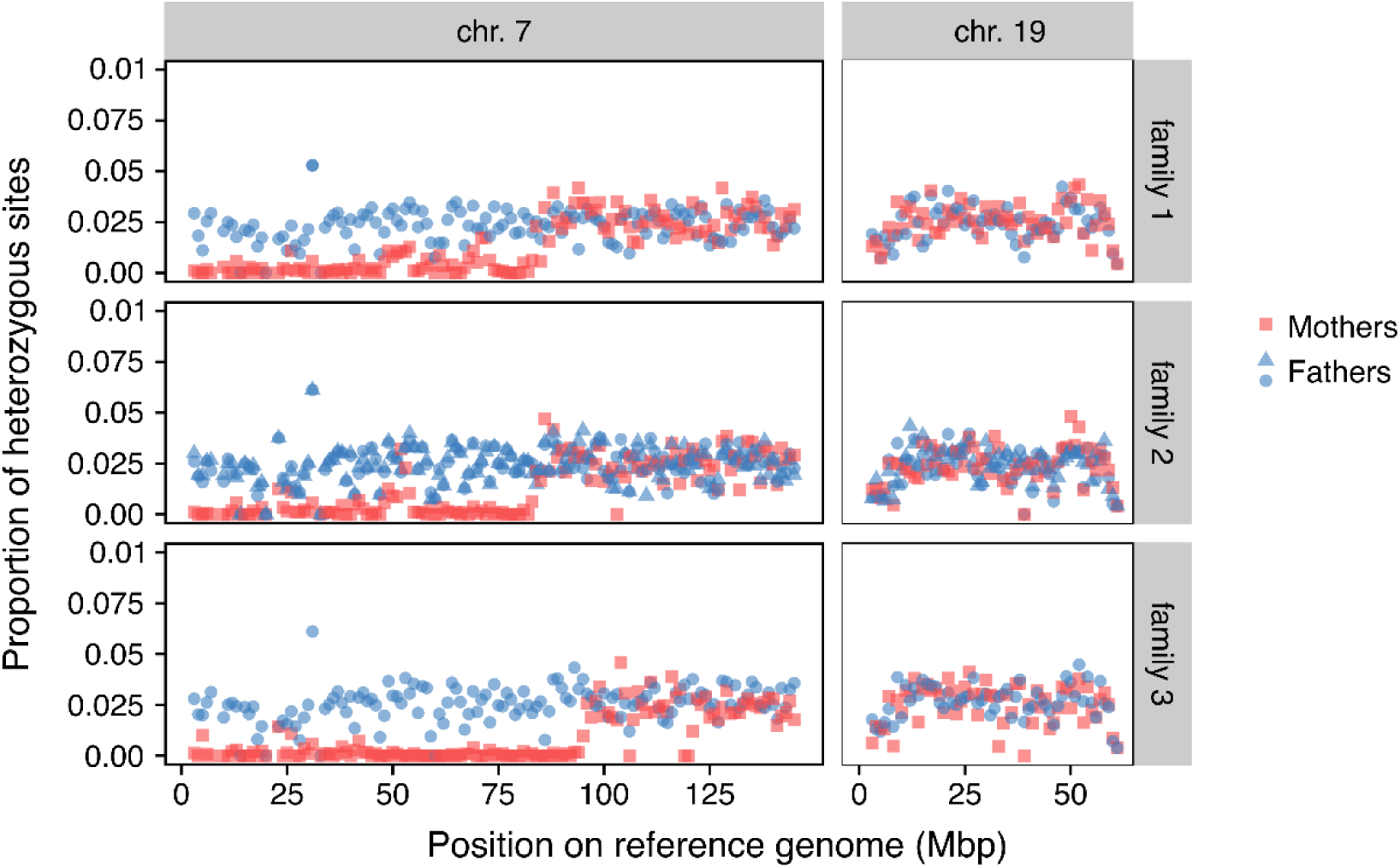
Proportion of heterozygous sites along the neo-sex chromosomes in mothers (red squares) and fathers (blue circles) based on the alignment position of RADtags to the *M. musculus domesticus* reference genome. Two fathers sired the offspring in family 2 and the heterozygosity for each is displayed (triangles instead of circles are used for one of the two fathers). Each point indicates the proportion of heterozygous sites averaged across a region of 1 megabase pairs (Mbp).

**Figure S7.**
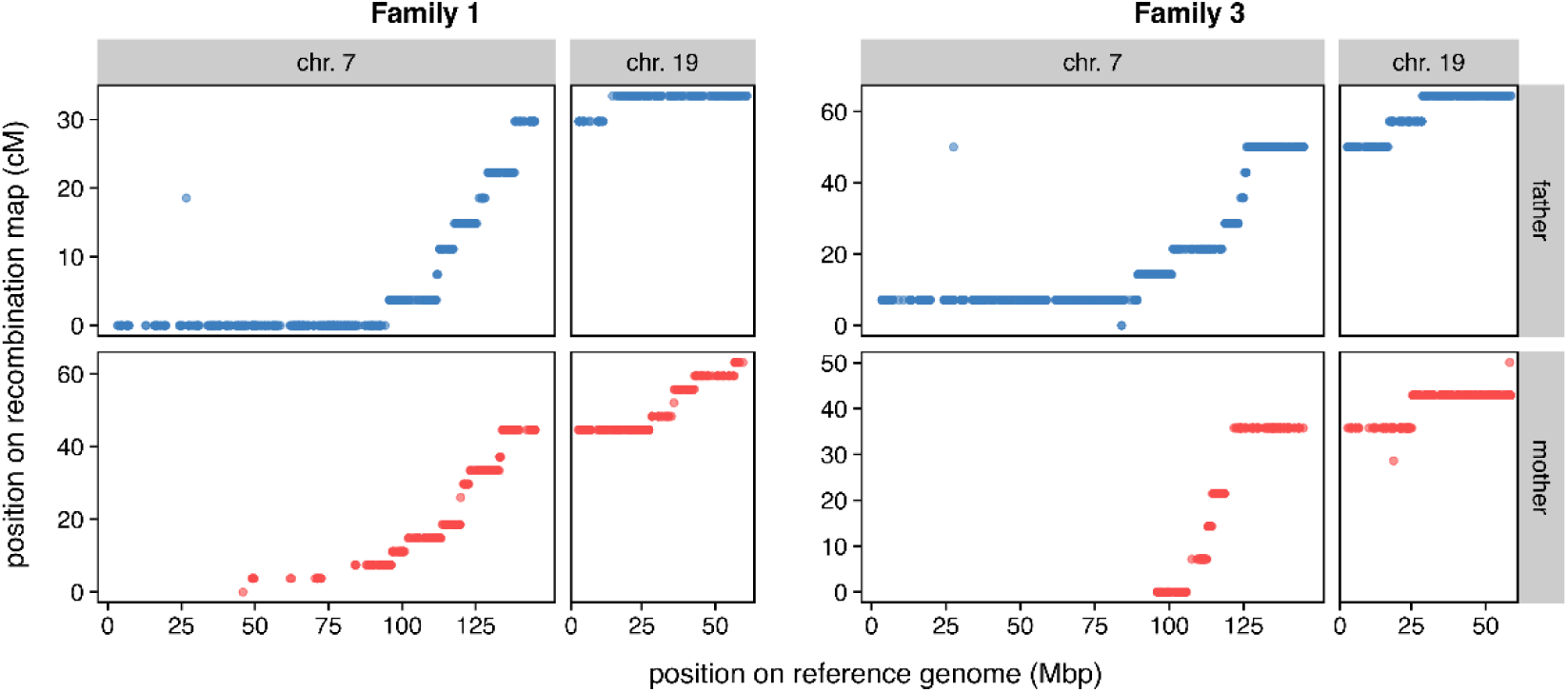
Recombination map of the fully sex-linked markers on the neo-sex chromosomes plotted against their alignment positions in the *M. musculus domesticus* reference genome. Each dot is a single marker (SNP); markers along the same horizontal line have segregated together within families. Breaks between lines of markers correspond to recombination events. cM: centi-Morgan, Mbp: megabase pairs.

**Table S1.** Number of fully sex-linked tags that align to the different chromosomes of the *Mus musculus domesticus* reference genome.

| Chromosome assigned | Male-limited tags | Fully sex-linked segregation pattern |  |  |
| --- | --- | --- | --- | --- |
|  |  | Family 1 | Family 2 | Family 3 |
| Chr 1 | 6 | 0 | 0 | 1 |
| Chr 2 | 0 | 0 | 0 | 1 |
| Chr 3 | 1 | 0 | 0 | 0 |
| Chr 4 | 0 | 0 | 0 | 0 |
| Chr 5 | 2 | 0 | 0 | 0 |
| Chr 6 | 0 | 0 | 0 | 0 |
| Chr 7 | 488 | 280 | 389 | 431 |
| Chr 8 | 0 | 0 | 1 | 0 |
| Chr 9 | 1 | 0 | 0 | 0 |
| Chr 10 | 1 | 0 | 1 | 0 |
| Chr 11 | 1 | 0 | 0 | 0 |
| Chr 12 | 0 | 0 | 1 | 0 |
| Chr 13 | 0 | 0 | 1 | 1 |
| Chr 14 | 2 | 0 | 0 | 0 |
| Chr 15 | 0 | 1 | 25 | 1 |
| Chr 16 | 0 | 0 | 0 | 0 |
| Chr 17 | 9 | 3 | 3 | 4 |
| Chr 18 | 1 | 0 | 0 | 0 |
| Chr 19 | 0 | 0 | 0 | 0 |
| Chr X | 3 | 0 | 0 | 0 |
| Chr Y | 23 | 0 | 0 | 0 |

**Table S2.** Alignment position of SNPs surrounding crossover events. The two SNPs shown for each crossover event (SNP “before” and “after”) are the closest ones found on two adjacent blocks on the recombination map (a block is defined as group of SNPs with the same genetic position on the recombination map).

| family | parent | crossover event | ID SNP before | ID SNP after | Chr. | Position of SNP before | Position of SNP after |
| --- | --- | --- | --- | --- | --- | --- | --- |
| family 1 | father | 1 | 76866_55 | 130733_49 | chr. 7 | 94267227 | 95593050 |
|  |  | 2 | 402771_86 | 195616_16 | chr. 7 | 111620423 | 111830934 |
|  |  | 3 | 238374_45 | 299543_65 | chr. 7 | 112060725 | 112663519 |
|  |  | 4 | 58200_82 | 57621_68 | chr. 7 | 117399976 | 117772451 |
|  |  | 5 | 306623_26 | 39768_21 | chr. 7 | 125305058 | 126174001 |
|  |  | 6 | 115891_85 | 186774_29 | chr. 7 | 128152403 | 129072804 |
|  |  | 7 | 279805_47 | 173462_85 | chr. 7 | 138160111 | 138392449 |
|  |  | 8 | 279805_47 | 173462_85 | chr. 7 | 138160111 | 138392449 |
|  |  | 9 | 165785_82 | 71316_15 | chr. 19 | 11839116 | 15037389 |
|  | mother | 1 | 90557_53 | 214335_68 | chr. 7 | 45923272 | 49015216 |
|  |  | 2 | 296068_47 | 298846_79 | chr. 7 | 72479031 | 83980649 |
|  |  | 3 | 81863_73 | 33406_91 | chr. 7 | 96225336 | 96681494 |
|  |  | 4 | 58314_42 | 157287_23 | chr. 7 | 100756993 | 102097635 |
|  |  | 5 | 140160_17 | 255732_30 | chr. 7 | 113142000 | 113615274 |
|  |  | 6 | 33468_9 | 260678_65 | chr. 7 | 119663035 | 119874621 |
|  |  | 7 | 33468_9 | 260678_65 | chr. 7 | 119663035 | 119874621 |
|  |  | 8 | 260678_65 | 11594_86 | chr. 7 | 119874621 | 120859858 |
|  |  | 9 | 361646_58 | 138162_74 | chr. 7 | 122609268 | 123097779 |
|  |  | 10 | 98115_23 | 277855_9 | chr. 7 | 132992466 | 133172673 |
|  |  | 11 | 406646_44 | 324769_41 | chr. 7 | 133398884 | 133980444 |
|  |  | 12 | 406646_44 | 324769_41 | chr. 7 | 133398884 | 133980444 |
|  |  | 13 | 30425_36 | 199542_74 | chr. 19 | 27446119 | 28451752 |
|  |  | 14 | 258291_86 | 164781_63 | chr. 19 | 35036993 | 35919361 |
|  |  | 15 | 164781_63 | 129528_59 | chr. 19 | 35919361 | 35980880 |
|  |  | 16 | 306422_40 | 415955_65 | chr. 19 | 42733271 | 42951393 |
|  |  | 17 | 388364_47 | 53367_22 | chr. 19 | 56336430 | 56574253 |
| family 3 | father | 1 | 243032_68 | 68284_67 | chr. 7 | 89217700 | 89457195 |
|  |  | 2 | 211306_89 | 33405_74 | chr. 7 | 100834226 | 101252283 |
|  |  | 3 | 10861_72 | 66449_49 | chr. 7 | 117703135 | 118755515 |
|  |  | 4 | 91267_69 | 14291_61 | chr. 7 | 123232373 | 124050039 |
|  |  | 5 | 276019_72 | 60419_12 | chr. 7 | 124994082 | 125333408 |
|  |  | 6 | 66955_39 | 156330_48 | chr. 7 | 125851650 | 125986888 |
|  |  | 7 | 457475_87 | 409111_35 | chr. 19 | 17176848 | 17625803 |
|  |  | 8 | 57623_58 | 441289_16 | chr. 19 | 28492391 | 28817150 |
|  | mother | 1 | 129436_27 | 336829_8 | chr. 7 | 105752333 | 107574345 |
|  |  | 2 | 194601_71 | 160780_34 | chr. 7 | 112682924 | 112967360 |
|  |  | 3 | 109550_43 | 46811_32 | chr. 7 | 114250716 | 114509038 |
|  |  | 4 | 371843_33 | 303262_88 | chr. 7 | 118623099 | 121643569 |
|  |  | 5 | 371843_33 | 303262_88 | chr. 7 | 118623099 | 121643569 |
|  |  | 6 | 100896_13 | 374702_53 | chr. 19 | 25108880 | 25439295 |

